# Redefining colorectal cancer classification and clinical stratification through a single-cell atlas

**DOI:** 10.1101/2021.02.02.429256

**Authors:** Ateeq M. Khaliq, Zeyneb Kurt, Miles W. Grunvald, Cihat Erdogan, Sevgi S. Turgut, Tim Rand, Sonal Khare, Jeffrey A. Borgia, Dana M. Hayden, Sam G. Pappas, Henry R. Govekar, Anuradha R. Bhama, Ajaypal Singh, Richard A. Jacobson, Audrey E. Kam, Andrew Zloza, Jochen Reiser, Daniel V. Catenacci, Kiran Turaga, Milan Radovich, Vineet Gupta, Ram Al-Sabti, Sheeno Thyparambil, Mia A. Levy, Janakiraman Subramanian, Timothy M. Kuzel, Anguraj Sadanandam, Arif Hussain, Bassel El-Rayes, Ameen A. Salahudeen, Ashiq Masood

**Author notes:** These authors contributed equally: Ateeq M. Khaliq, Zeyneb Kurt and Miles W. Grunvald.

## Abstract

Colorectal cancer (CRC), a disease of high incidence and mortality, has had few treatment advances owing to a large degree of inter- and intratumoral heterogeneity. Attempts to classify subtypes of colorectal cancer to develop treatment strategies has been attempted by Consensus Molecular Subtypes (CMS) classification. However, the cellular etiology of CMS classification is incompletely understood and controversial. Here, we generated and analyzed a single-cell transcriptome atlas of 49,859 CRC cells from 16 patients, validated with an additional 31,383 cells from an independent CRC patient cohort. We describe subclonal transcriptomic heterogeneity of CRC tumor epithelial cells, as well as discrete stromal populations of cancer-associated fibroblasts (CAFs). Within CRC CAFs, we identify the transcriptional signature of specific subtypes (CAF-S1 and CAF-S4) in more than 1,500 CRC patients using bulk transcriptomic data that significantly stratifies overall survival in multiple independent cohorts. We also uncovered two CAF-S1 subpopulations, ecm-myCAF and TGFß-myCAF, known to be associated with primary resistance to immunotherapies. We demonstrate that scRNA analysis of malignant, stromal, and immune cells exhibit a more complex picture than portrayed by bulk transcriptomic-based Consensus Molecular Subtypes (CMS) classification. By demonstrating an abundant degree of heterogeneity amongst these cell types, our work shows that CRC is best represented in a transcriptomic continuum crossing traditional classification systems boundaries. Overall, this CRC cell map provides a framework to re-evaluate CRC tumor biology with implications for clinical trial design and therapeutic development.

## INTRODUCTION

Colorectal cancer (CRC) is the third most commonly diagnosed cancer and a leading cause of cancer-related mortality worldwide^1, 2^. Approximately one-third of patients experience disease relapse following curative-intent surgical resection and chemotherapy^3^. Despite the high incidence and mortality of advanced CRC, few effective therapies have been approved in the past several decades^4^. One barrier to the development of efficacious therapeutics is the biological heterogeneity of CRC and its variable clinical course. While landmark studies from The Cancer Genome Atlas (TCGA) have defined the somatic mutational landscape within CRC, several studies have shown that stromal signatures, including fibroblasts and cytotoxic T cells, are likely the main drivers of clinical outcomes^5–9^. These findings suggest that the clinical phenotypes of CRC and by extension, its tumor biology is shaped by a complex niche of heterotypic cell interactions within the tumor microenvironment (TME).

Bulk gene expression analyses by several independent groups have identified distinct CRC subtypes^10–12^. Based on bulk transcriptomic signatures, an international consortium published the Consensus Molecular Subtypes (CMS) that classified CRC into CMS1 (MSI immune), CMS2 (canonical), CMS3 (metabolic), and CMS4 (mesenchymal) subtypes^12^. Unfortunately, associations between CMS and meaningful therapeutic responses to specific agents have been inconsistent across studies^13–16^. Further, there has been a lack of concordance between primary and metastatic CRC tumors within the CMS framework, limiting its overall utility in clinical decision making^16–18^. Thus, an improved CMS classification, or an alternative classification system, is needed to improve clinical utility.

To overcome the limitations of bulk-RNA sequence profiling, we utilized single-cell RNA sequencing (scRNA-seq) to more thoroughly evaluate the CRC subtypes at the molecular level, including within the context of the currently defined CMS classification. We dissected heterotopic cell states of tumor epithelia and stromal cells, including a cancer-associated fibroblast (CAF) population. The CAF population’s clinical and prognostic significance became apparent when CAF signatures were applied to large, independent CRC transcriptomic cohorts.

## RESULTS

We profiled sixteen primary CRC tumor tissue samples and eight adjacent, normal, colonic tissue samples (24 in total) using droplet-based, scRNA-seq. Altogether, we captured and retained 49,589 high-quality single cells after performing quality control for downstream analysis (Fig. 1a, **Supplementary Table 1**). All scRNA-seq data were merged and normalized to identify robust discrete clusters of epithelial cells (*EPCAM+, KRT8+,* and *KRT18+*), fibroblasts (*COL1A2+*), endothelial cells (*CD31+*), T cells (*CD3D+),* B cells *(CD79A+),* and myeloid cells (*LYZ+*) using canonical marker genes (Fig. 1b-c). Additionally, each cell type compartment was analyzed separately. Clustree (v0.4.1) and manual review of differentially expressed genes in each subcluster were studied to choose the best cluster resolution without cluster destabilization (see methods)^19^. Cell population designation was chosen by specific gene expression, and SingleR was also utilized for unbiased cell type recognition (see methods)^20–23^.

**Figure 1.**
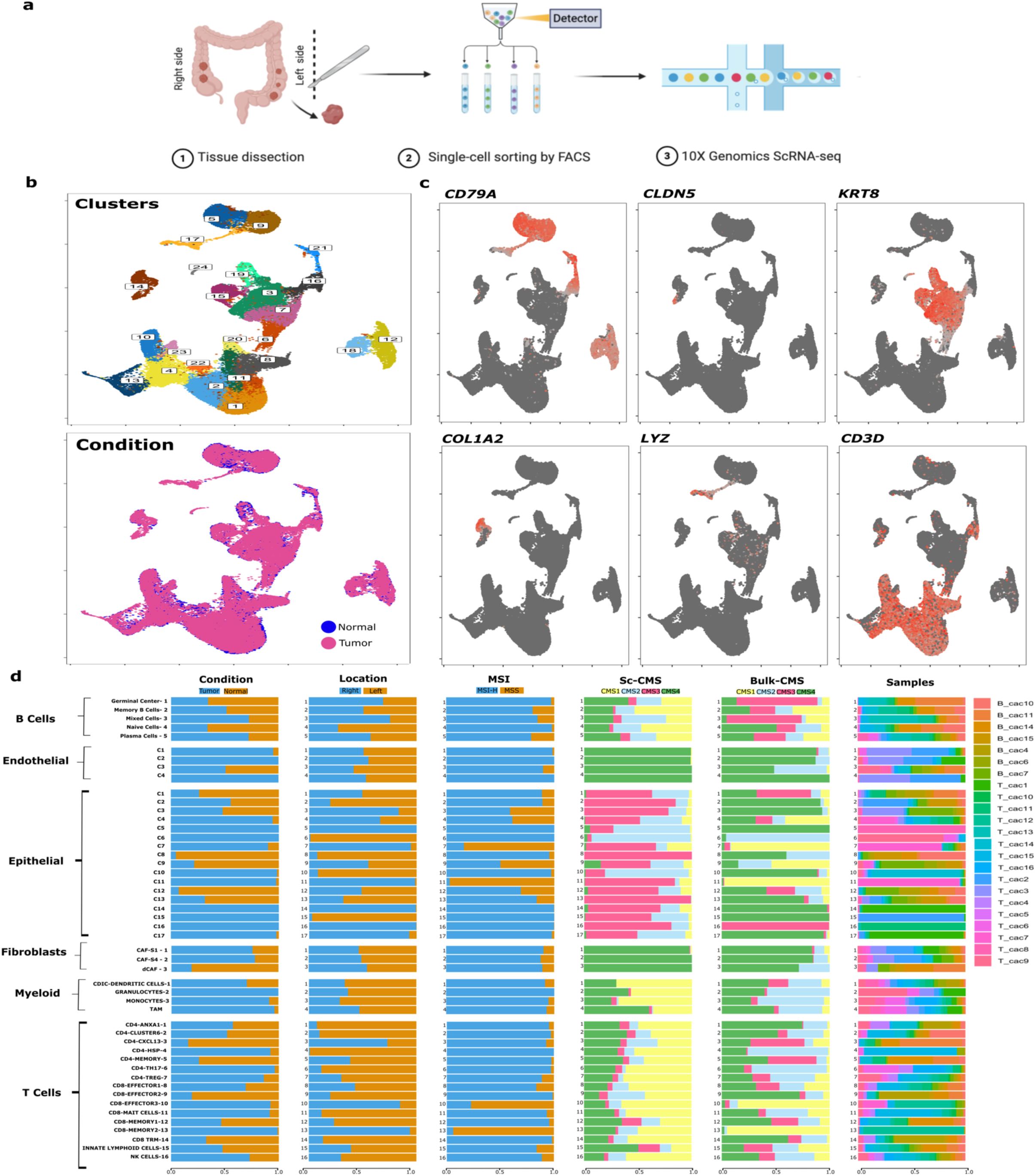
Identification and clustering of single cells. **a,** Workflow of sample collection, sorting, and sequencing (methods contain full description for each step). **b,** UMAP characterization of the 49,859 cells profiled. Coloring demonstrates clusters, tumor vs. normal sample origin (condition), and individual sample origin. **c,** Identification of various cell types based on expression of specified marker genes. **d,** Characterization of the proportion of cell types identified in tumor vs. normal colon tissue, sidedness (right vs. left), microsatellite instability (MSI) status, single-cell Consensus Molecular Subtypes (scCMS) classification, Consensus Molecular Subtypes (CMS) of bulk RNA-seq data, and origin of sample. The transcription counts of tumor and normal tissue cell types are demonstrated at the bottom with boxplot representation. The graph represents total clusters and cell types identified after re-clustering of each cell compartment depicting global heterogeneous landscape of colorectal cancers.

In addition to cancer cells, we identified diverse TME cell phenotypes, including fibroblasts subsets (*CAF-S1* and *CAF-S4*), endothelial cells, CD4+ subsets (naïve/memory, Th17, and Tregs), CD8+ subsets (naïve/memory, cytotoxic, tissue-resident memory, and Mucosa-Associated Invariant (MAIT) cells), NK cells, innate lymphoid cell (ILC) types, B cell phenotypes (naïve, memory, germinal center, and plasma cells), and monocyte lineage phenotypes (C1DC+ dendritic cells, proinflammatory monocytes [IL1B, IL6, S100A8, and S100A9]), and M2 polarized anti-inflammatory [*CD163, SEPP1, APOE,* and *MAF]*), tumor-associated macrophages (TAMs) (Fig. 1d, Extended Data Figs. 1-3, **Extended Data Tables 1-4**)^20–22, 24^.

We also profiled an independent CRC dataset for validation and retained 31,863 cells after quality control and strict filtering of cells expressing hybrid markers^25^. The re-clustering of individual compartments further refined our analysis, and cells that expressed hybrid or distinct lineage markers within a cluster were removed from the downstream analysis (Supplementary Fig. 4-5, 9-10) (see methods). Thus, a total of 81,242 high-quality cells were profiled to produce a single-cell map of 39 colorectal cancer patients. The results of the primary CRC cohort (49,859 single-cells) are available at the Colon Cancer Atlas (www.websiteinprogress.com).

### Malignant colon cancer reveals tumor epithelial cell subclonal heterogeneity and stochastic behavior

We detected 8,965 tumor and benign epithelial cells (*EPCAM+, KRT8+,* and *KRT18+*) and, on re-clustering, produced 17 epithelial clusters (designated C1 to C17) (Fig. 2a-b). Tumor cells were confirmed to be of malignant origin by inferring chromosomal copy number alterations (Supplementary Fig. 1). Clusters were chiefly influenced by colonic epithelial markers, including those for stemness (*LGR5, ASCL2, OLFM4,* and *STMN1*), enterocytes (*FABP1* and *CA2*), goblet cells (*ZG16, MUC2, SPINK4,* and *TFF3*), and enteroendocrine cells (*PYY* and *CHGA*) (**Supplementary Table 2**). Tumor cells exhibited a high degree of de-differentiated state of plasticity, possibly accounting for lasting cancer growth (Supplementary Fig. 2b)^26^. Distinct tumor-derived clusters were predominantly patient-specific, reflecting a high degree of inter-patient tumoral cell heterogeneity (Fig. 1d). In contrast, epithelial populations derived from normal colon tissue samples across multiple patients clustered together, a pattern observed in previous studies confirming both normal tissue homeostasis and limited sample batch effects (Fig. 1d)^27, 28^.

**Figure 2.**
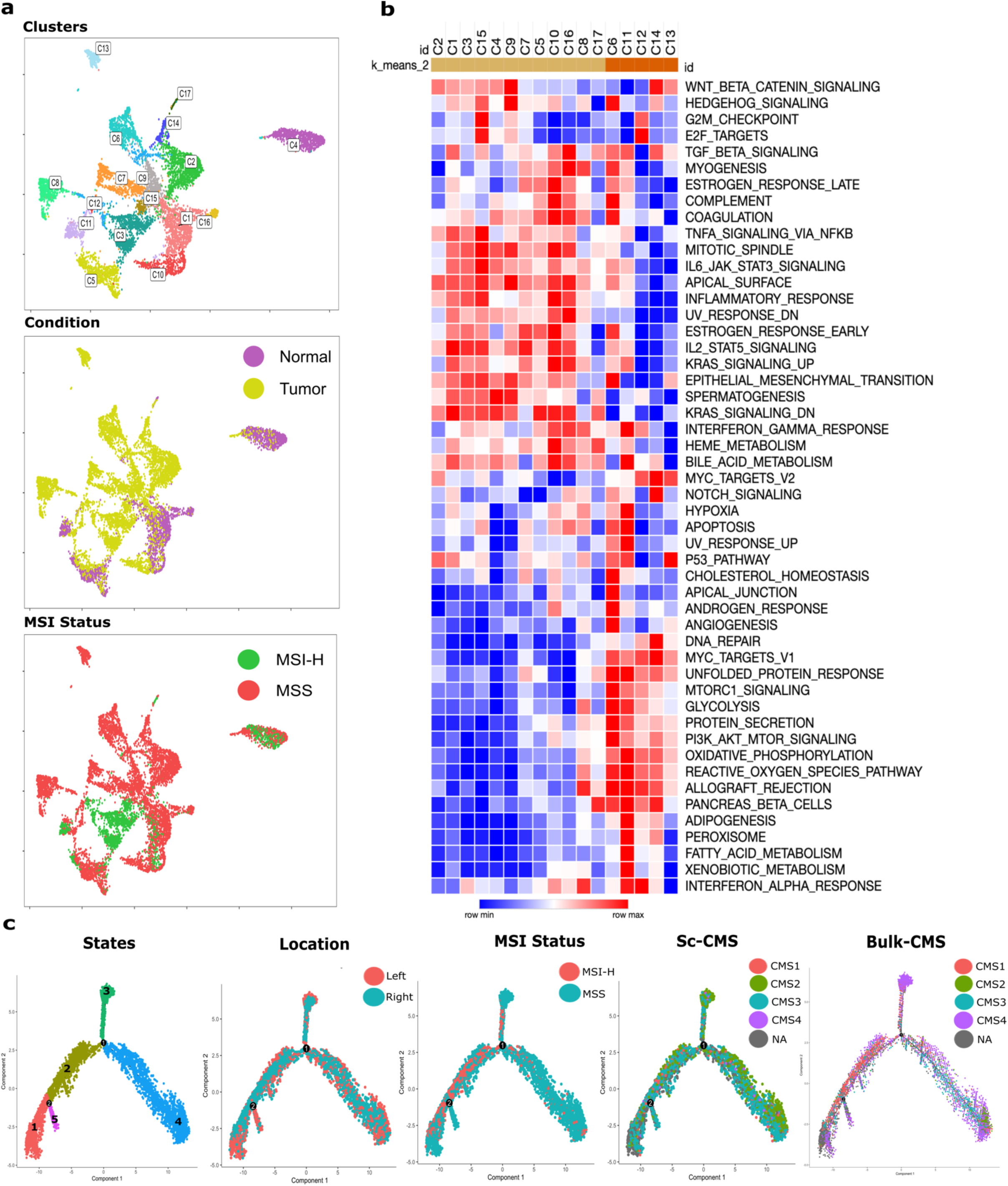
Reclustering and characterization of the epithelial compartment. **a**, UMAP of tumor and non-malignant epithelial reclustering demonstrating 17 distinct clusters. **b,** Bar plot representation of cell proportions by sample, tissue type, MSI status, tumor location, scCMS score, and bulk CMS score. **c,** Heatmap of Hallmark pathway analysis within the epithelial cell compartment. **d,** Trajectory analysis of cells colored by tumor location, scCMS, MSI status, and bulk CMS status.

We next aimed to identify gene expression programs shared across these clusters using hallmark pathway analysis^29^. A strong overlap was observed for multiple pathways such as activation of inflammatory, epithelial-mesenchymal transformation (EMT), immune response, and metabolic pathways (Fig. 2b). Interestingly, high microsatellite instability (MSI-H) and microsatellite stable (MSS) CRC tumors, considered clinically separate entities, demonstrated similar pathway program activation within the tumor epithelial populations (Fig. 2b). Some clusters also showed activation of unique pathways such as activation of apical junctions and angiogenesis (C6), hypoxia and fatty acid metabolism (C11) and Notch signaling and DNA repair (C14), among others (Fig. 2b). However, MSI-H tumors differed from MSS tumors based on immune cell infiltration (Extended Data Fig. 1).

Since intratumoral heterogeneity is recognized as a key mechanism contributing to drug resistance, cancer progression, and recurrence, we next focused on dissecting potential transcriptomic states to identify heterogeneity within each tumor^30–32^. We found that each tumor specimen contained 2-10 distinct tumor epithelial clusters (Fig. 1d). Gene set variation analysis (GSVA) was performed on cells from individual tumor samples and illustrated the sub-clonal transcriptomic heterogeneity within each specimen (Supplementary Fig. 2c)^33^. Clusters identified in individual pathway analysis demonstrated the up-or down-regulation of crucial metabolic and oncogenic pathways between samples, suggesting wide phenotype variations between cells from the same tumor^34^.

Given the evidence of intratumoral epithelial heterogeneity, we next performed trajectory inference using pseudotime analysis to identify potential alignments or lineage relationships (i.e., right versus left-sided CRC), CMS classification, or MSI status^35, 36^. This analysis also served as a control for inter-patient genomic heterogeneity and provided an orthogonal strategy to confirm the transcriptomic trends we identified. We detected five molecular states (S1n/t to S5n/t) with malignant and normal epithelial cells intermixed and aligned along a common transcriptional trajectory (Fig. 2c)^37, 38^. In both normal and tumor cells, each transcriptional state pathway activation was shared and related to colon epithelial function of nutrient absorption, maintaining the colon homeostasis, and the activation of cancer-related pathways such as apoptosis and cell development^39^. However, these cell states showed differential gene enrichment activation between normal and tumor cells (Supplementary Fig. 3**, Supplementary Table 3)**.

Additionally, tumor cells showed upregulation of embryogenesis (S2t), consistent with previous findings that tumor cells revert to their embryological states in cancer development (**Supplementary Table 3**)^40^. Interestingly, there were no significant associations with anatomic location, CMS classification, or MSI status within our dataset or an independent dataset of 31,383 single cells (Fig. 2d, Supplementary Fig. 4)^25^. Hence, in our analysis, CRC oncogenesis represent the hijacking of the normal epithelial differentiation pathways coupled with the acquisition of embryonic pathways^37, 40^.

### CRC-associated fibroblasts in the tumor microenvironment exhibit diverse phenotypes, and specific subtypes are associated with poor prognosis

We next focused on CRC TME subpopulations. 819 high-quality fibroblasts were re-clustered into eight clusters, and then phenotypically classified into two major subtypes to assess for further CAF heterogeneity. These phenotypic subtypes were found to be immunomodulatory CAF-S1 (*PDGFRA+* and *PDPN+*) and contractile CAF-S4 (*RGS5+* and *MCAM+*) (Fig. 3a-b**, Supplementary Table 4**)^41, 42^. This fibroblast cluster dichotomy was also observed in the independent CRC patient scRNA-seq dataset of 31,383 cells (Supplementary Fig. 5)^25^.

**Figure 3.**
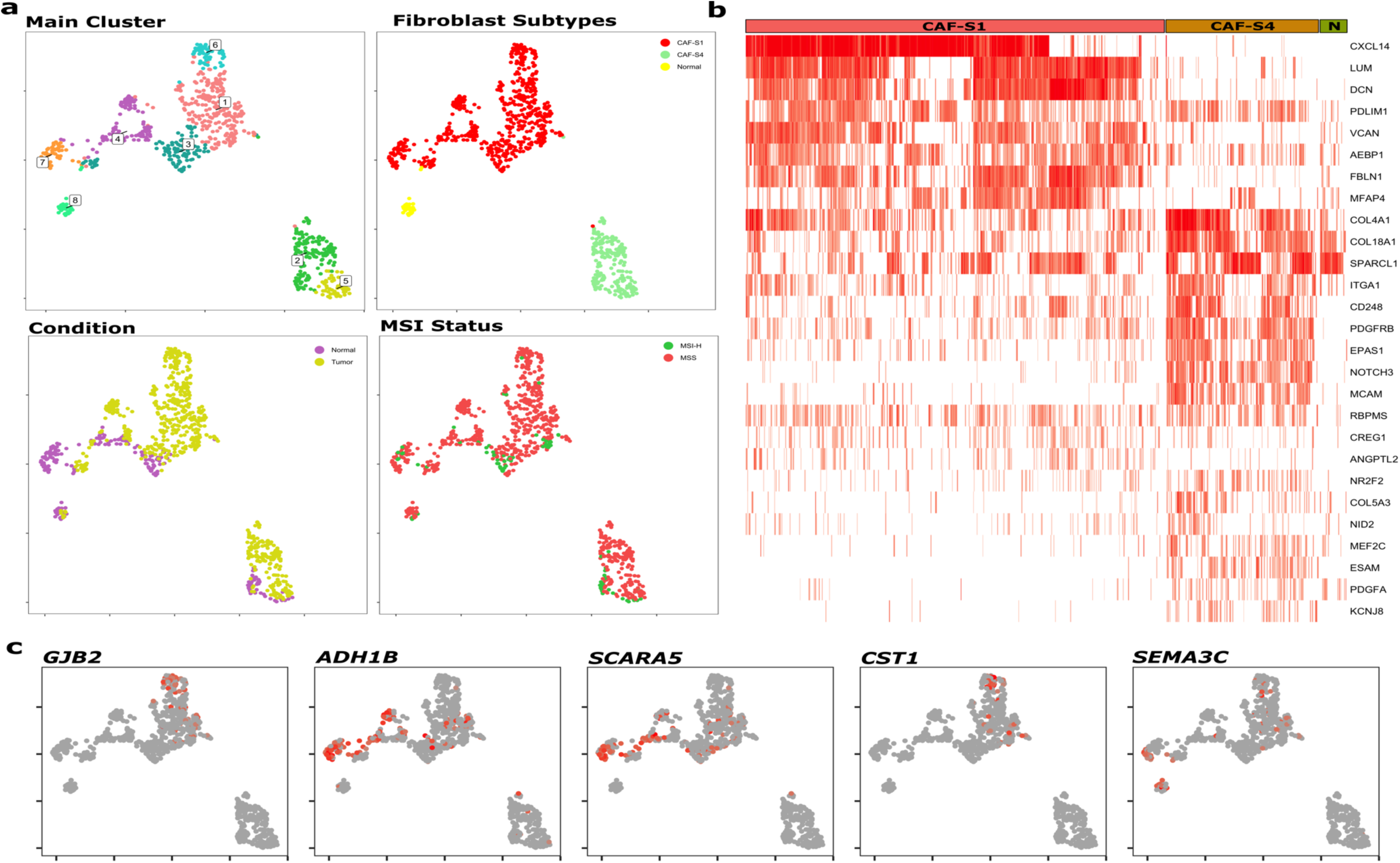
Fibroblast clusters in colon and colorectal tumors. **a**, UMAP of 819 fibroblasts colored by distinct clusters, CAF status, tissue status and origin of sample. UMAP of fibroblasts colored by specific CAF-S1 subtypes. **b,** Heatmap showing the variable expression of fibroblast specific marker genes across CAF-S1, CAF-S4, and normal fibroblasts. **c,** UMAP color-coded for marker genes for five CAF-S1 subtypes as indicated.

The CAF-S1 and CAF-S4 subtypes showed striking resemblances to the mCAF (extracellular matrix) and vCAF (vascular) fibroblast subtypes, respectively, as previously described in a mouse breast cancer model^43^. Most clusters were found in multiple patients, albeit in varying proportions, signifying shared patterns in CAF transcriptomic programs between patients. Fibroblasts derived from MSI-H tumors were distributed similarly throughout these clusters (Fig. 1d).

CAF-S1 cells exhibited high chemokine expressions such as *CXCL1, CXCL2, CXCL12, CXCL14,* and immunomodulatory molecules including *TNFRSF12A* (**Supplementary Table S4)**. Additionally, CAF-S1 cells expressed extracellular matrix genes including matrix-modifying enzymes (*LOXL1* and *LOX*)^43^. To determine this population’s functional significance, we compared the CAF-S1 population transcriptomes to those described recently in breast cancer, lung cancer, and head and neck cancer^44^. We recovered five CAF-S1 subtypes that included ecm-myCAF (extracellular; *GJB2*), IL-iCAF (growth factor, *TN*F and interleukin pathway; *SCARA5*), detox-iCAF (detoxification and inflammation; *ADH1B*), wound-myCAF (collagen fibrils and wound healing; *SEMA3C*), and TGFβ-myCAF (*TGF-β* signaling and matrisome; *CST1, TGFb1*), which were previously divided into two major subtypes: iCAF and myCAF (Fig. 3c). Among these five subtypes, ecm-myCAF and TGFβ-myCAF are known to correlate with immunosuppressive environments and are enriched in tumors with high regulatory T lymphocytes (Tregs) and depleted CD8+ lymphocytes. Additionally, these subtypes are associated with primary immunotherapy resistance in melanoma and lung cancer^44^.

The CAF-S4 population expressed pericyte markers (*RGS5+, CSPG4+,* and *PDGFRA+*), *CD248 (endosialin)*, and *EPAS1 (HIF2-α),* that this particular CAF subtype is vessel-associated, with hypoxia potentially contributing to invasion and metastasis, as has been shown in another study (Fig. 3a-b**, Supplementary Table 4)**^43^. CAF-S4 clustered into the immature phenotype (*RGS5+, PDGFRB+,* and *CD36+)* and the differentiated myogenic subtype (*TAGLN+* and *MYH11+*) **(Supplementary Table 4)**^42^.

Given the correlation between CMS4 and fibroblast infiltration, we next sought to test the existence of CAF-S1 and CAF-S4 signatures in bulk transcriptomic data and their association with clinical outcomes^12^. To this end, we interrogated and carried out a meta-analysis of eight colorectal cancer transcriptomic datasets comprising 1,584 samples and confirmed the presence of CAF-S1 and CAF-S4 gene signatures in CRC and other cancer types (Fig. 4a). We detected a strong and positive correlation between specific genes differentially expressed between each CAF subtype in CRC (Fig. 4a-d)^43^. We also confirmed the presence of CAF-S1 and CAF-S4 signatures in pancreatic adenocarcinoma (n=118) and non-small cell lung cancer (NSCLC, n=80) cohorts (Fig. 4a) (see methods for datasets). Gene signatures were specific to each CAF-S1 and CAF-S4 in bulk transcriptomic datasets thus confirming their existence in TME of CRC and other tumor types.

**Figure 4.**
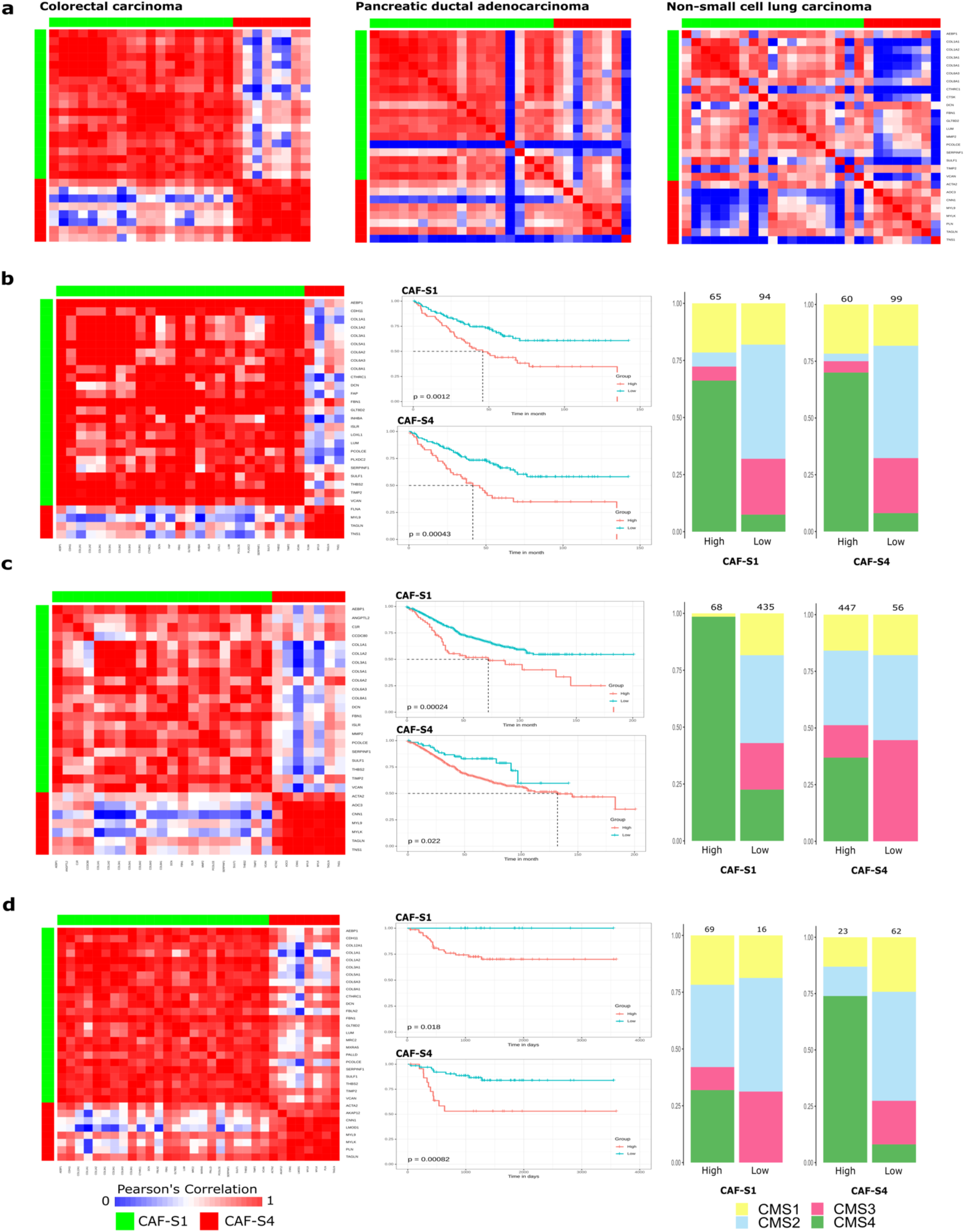
Correlation of CAF-S1 and CAF-S4 gene profiles across human bulk transcriptomic data. **a**, Pearson’s correlation of genes from CAF-S1 and CAF-S4 profiles in colorectal cancer (n= 1584; CAF-S1 and CAF-S4 r > 0.8), pancreatic cancer (n= 118; CAF-S1 r = 0.70; CAF-S4 r= 0.60), non-small cell lung cancer (n = 80; CAF-S1 r = 0.69; CAF-S4 r = 0.67). **b-d,** Pearson correlation plots, Kaplan-Meyer survival curves, and bar plots of CMS status assessing CAF expression in individual CRC datasets. Plots b-c are generated from single GEO datasets; GSE17536 (n = 177), GSE39582 (n= 585) and GSE33113 (n= 96), respectively. Note: High CAF-S1 and CAF-S4 gene signatures are associated with poor survival across all CMS . r = coefficient correlation. Hazard ratio > 1 in figures b-d.

We found high CAF-S1 and CAF-S4 signatures associated with significantly poor median overall-survival, irrespective of CMS in three independent CRC datasets (Fig. 4b-d, Supplementary Fig. 6). Additionally, CAF signatures stratified the CMS4 subtype into high-and low-risk overall survival in all datasets, thus identifying additional heterogeneity and providing prognostication in this aggressive patient subgroup (Fig. 4b-d). Here, using scRNA-seq, we show for the first time that high CAF infiltration in CRC is associated with poor prognosis across all molecular subtypes, and which further stratifies the CMS4 subgroup into high and low-risk clinical phenotypes in CRC cohorts.

### Single-cell RNA sequencing reveals heterogeneity beyond Consensus Molecular Subtypes in colorectal cancers and offers therapeutic opportunities

The lack of association between tumor epithelia and CMS classification, as well as the survival differences between high-and low-risk CAF signatures across CRC molecular subtypes suggest CRCs are less well defined than the traditional classification systems have indicated (e.g. those systems defined by somatic alterations, epigenomic features, and bulk gene expression data)^10–12, 45, 46^.

To test our hypothesis, we estimated every cell type fraction using single-cell data in a discovery cohort (GSE39852^11^, n=585) with a machine-learning algorithm, CIBERSORTx^47^. Cell type fractions were then validated in two independent cohorts of 177 (GSE17536^48^) and 290 (GSE14333^49^) tumors^9^. When we compared epithelial, immune, and stromal cell populations among the CMS subtypes, we did not detect a distinct pattern of tumor, immune, or stromal cell abundance across the four subtypes (Fig. 5a-b, Supplementary Figs. 7a-b, 8a-b). Although, we noted some trends in cell pattern enrichment amongst a few cell types consistent with CMS classification (CAFs were enriched in CMS4; dendritic cells, monocytes and TAMs were enriched in CMS1/CMS4; epithelial cells were enriched in CMS3) overall cellular phenotypes were present in varying proportions without a clear distinction between the four subtypes. The discovery and validation cohorts showed significant discordance in terms of cell phenotype enrichment with respect to each CMS subtype (Fig. 5a-b, Supplementary Figs. 7a-b, 8a-b)^9^. These discordant results could potentially be due to intra-tumoral variations in tumor purity, location of tumor biopsy, stromal and immune cell infiltration, and/or CMS’s inability to address tumor-TME to tumor-TME and TME to TME variabilities, among other factors^9, 50^.

**Figure 5.**
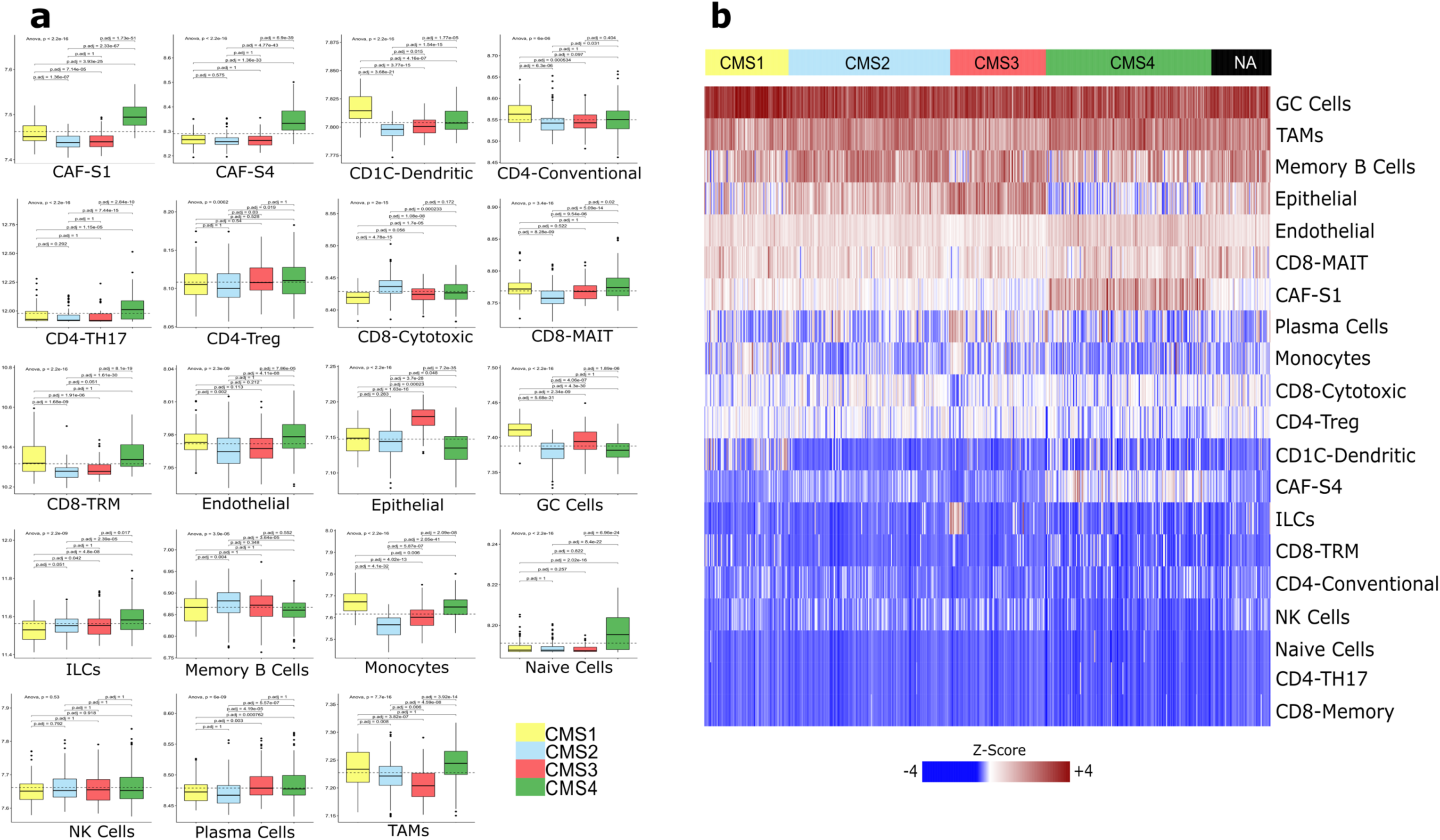
Average cell type abundance from CRC dataset GSE39582 and sorted by bulk CMS status. **a**, Boxplots show the distribution of cell types within tumors with varying CMS status. The whiskers depict the 1.5 x IQR. The p-values for pairwise t-tests comparisons (with Benjamani-Hochberg correction) and ANOVA tests of cell abundance across CMS are shown in the figure. **b,** Deconvolution heatmap of different cell types by average expression using CIBERSORTx demonstrating cell type distribution within each CMS category. ILCs = Innate lymphoid cells, GC Cells = Germinal Center B Cells, NK Cells = Natural Killer Cells, TAMs = Tumor Associated Macrophages.

Based on the above findings, we hypothesized that CRC tumors may be more accurately represented as a continuum as has been proposed by Ma et al.^51^. The authors analyzed bulk transcriptomic data using a novel computational framework in which *denovo*, unsupervised clustering methods (k-medoid, non-negative-matrix factorization, and consensus clustering) best classified CRC tumors in a transcriptomic continuum^52–54^. They further carried out principal component analysis and robustly validated two principal components, PC Cluster Subtype Scores 1 and 2 (PCSS1 and PCSS2, respectively). Using this framework, we reasoned that single-cell data could elucidate the biological underpinnings of a CRC continuum model, and resolve stromal confounding seen using bulk transcriptomes^50, 55^.

We evaluated every cell fraction (epithelial, stromal, and immune components) using the Ma et al. algorithm on our discovery and validation cohorts, focusing on the validated PCSS1 and PCSS2. Upon projecting bulk transcriptomes onto the four CMS quadrants, we analyzed single cell-specific gene signatures after deconvolution. Intriguingly, we noted that each cell type not only projected in the expected quadrant but some of these same cell types also projected on other CMS quadrants (Fig. 6a-b**, supplementary 9a-b, 10a-b, Supplementary table 5-6)**. For example, epithelial cells, in addition to projecting on CMS2/CMS3 quadrants, also projected on other CMS quadrants and exhibited continuum. These data reflect significant intra-tumoral heterogeneity among all the cellular components that makeup the tumor ecosystem. Thus, it appears that CRC exists in a transcriptomic continuum not only with respect to the tumor cells themselves but also all the other cell types that make up the TME. These aspects would not have been apparent based on bulk transcriptomics analysis alone.

**Figure 6.**
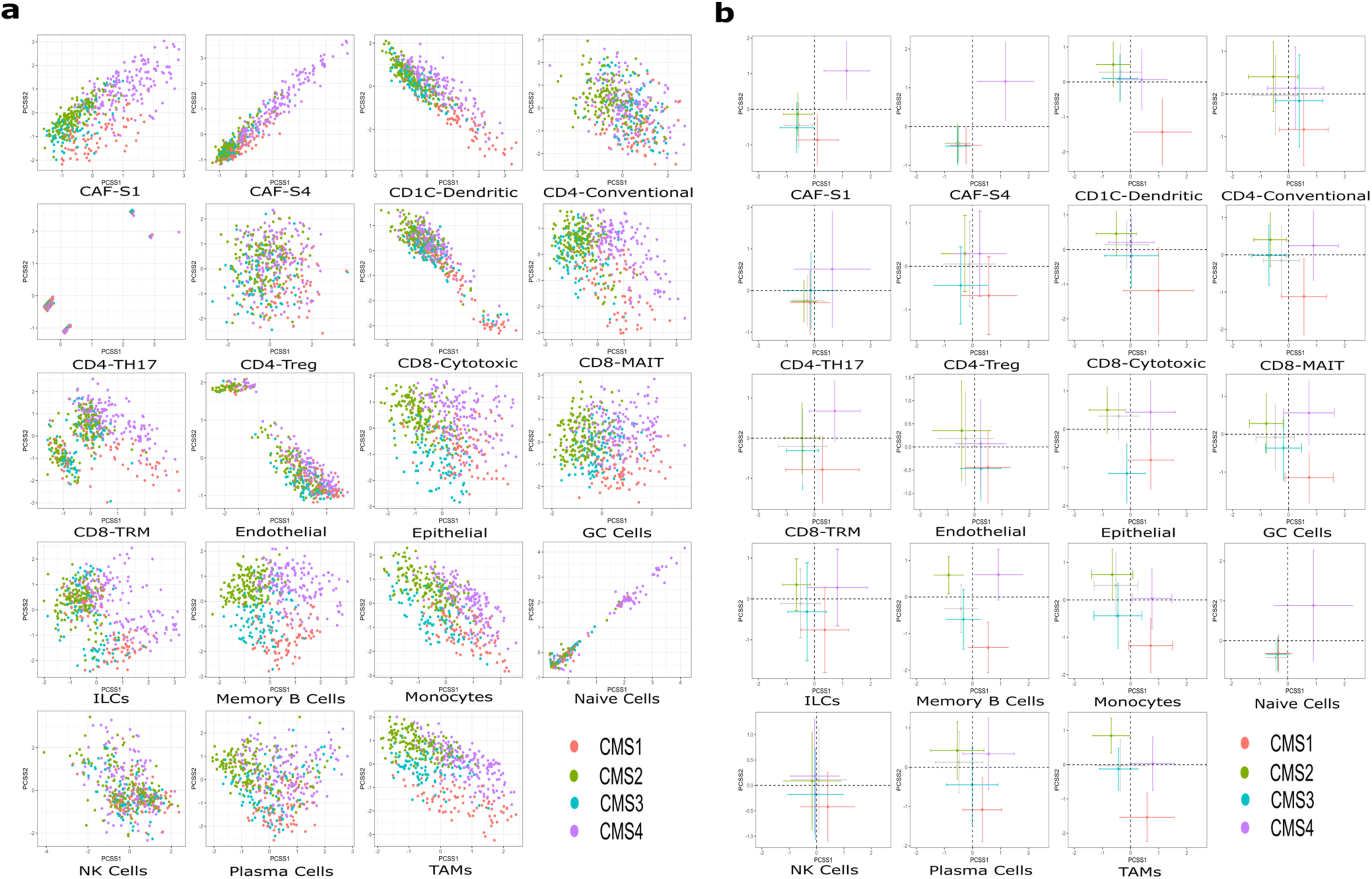
Continuous scores for CRC dataset GSE39582. **a,** Principal component analysis plot showing PCSS1 and PCSS2 continuous scores reported by CMS classification across 19 cell types show minimal separation in the top 2 principal components. **b,** All cell types projected on four quadrants representing CMS1-4 using PCSS1 and PCSS2 scores. Note that the cell types largely form a continuum along CMS status and are not clustered in discrete quadrants separate from one another. Cells and markers are colored by bulk CMS status accordingly to the tumor sample of origin. ILCs = Innate lymphoid cells, GC Cells = Germinal Center B Cells, NK Cells = Natural Killer Cells, TAMs = Tumor Associated Macrophages.

We found that transcriptional shifts were reproducible across discovery and validation datasets for major cell types (Fig. 6a-b**, supplementary 9a-b, 10a-b)**. Our analysis show no reliability in classifying CRC into immune–stromal rich (CMS1/CMS4) or immune-stromal desert (CMS2/CMS3) subtypes as proposed previously^16^. Thus, confirming continuous scores rather than discrete subtypes may improve classifying CRC tumors and may explain tumor-to-tumor variability, tumor/TME-to-tumor/TME and TME-to-TME variabilities within CMS subgroups as has been previously described ^51^. While CAF-S1 and CAF-S4 were present across all CMS groups these CAFs exhibited high PCSS1 and PCSS2 scores correlating with increased enrichment in the CMS4 group. Thus, our analysis identified CAF-S1 and CAF-S4 as the cells of origin for biological heterogeneity in CMS4 subtypes associated with poor prognosis (**Supplementary table 5-6**).

## DISCUSSION

In the present study, we evaluated the CMS classification system of CRC that has been established using bulk RNA-seq through the lens of single-cell transcriptomics. We identified significant intratumoral transcriptomic heterogeneity of cell states within tumor epithelial cells. These findings are consistent with a prior DNA-based multi-omics approach by Sottoriva et al. which proposed a ‘big bang model’ of CRC^56^. This model suggests that at initiation, CRC is composed of a mixture of subclones, underscoring the significant heterogeneity inherent in CRC biology and potentially reflecting non-linear pathways during CRC evolution and progression. Furthermore, both our findings (using scRNA-seq and pseudotime trajectory analysis) and Sottoriva et al. agree with clinical observations that tumors with diverse clonal populations respond unpredictably to monolithic treatment strategies “targeted” to average expression profiles based on bulk sequencing. Thus, targeting sub-clonal transcriptomic patterns is likely to be more effective when designing personalized CRC therapies ^56 57^. Interestingly, a recent study in gastroesophageal adenocarcinoma (GEA) that employed therapies to target dominant sub-clonal populations in metastatic GEA patients demonstrated improved survival as compared to standard monolithic treatment strategies^58^.

We find that stromal cells are important contributors to the observed biological heterogeneity of CRC, in agreement with emerging data that suggest stromal-derived signatures play key role in modulating CRC biology^7–9, 37, 59, 60^. These recent studies employing bulk transcriptomics demonstrated that the degree of stromal infiltration is associated with prognosis, while a small scRNAseq study utilizing 26 fibroblasts showed poor survival particularly among the CMS4 subtype CAF-enriched CRC tumors^61^. By using a much larger sample set of 1,182 high-quality fibroblasts, our current work identifies the CAF-S1 and CAF-S4 subtypes to be the cells of origin associated with poor prognosis across all CRC patients and not just those with CMS4 classified tumors^61^. CAF signatures are able to stratify patients by median overall survival (Fig. 4b-d). These findings are significant since the CMS4 subtype, is primarily stromal-driven and is enriched in more than 40% of metastatic CRC samples from patients with worse outcomes^17, 18^. Thus, targeting CAFs to remodel the tumor microenvironment may lead to improved and much-needed therapeutic development for metastatic CRC patients^17, 18^.

Targeting of CAFs in solid tumors is being explored in multiple clinical trials with variable results^62, 63^. Such studies likely failed to address CAF heterogeneity and their complex interactions with the other cells of TME. Our study suggests that CRC may be intricately entwined with the stroma, and therefore may be amenable to stromal targeted combinatoric approaches, including monoclonal antibodies that abrogate CAF-S1 function. In future studies, the treatment of CRC patients should involve stroma targeted therapies and take the above aspects into consideration^63, 64^. The scRNA-seq or bulk-RNA-seq signatures corresponding to CAF-S1 and CAF-S4 may serve as suitable biomarkers for tumors that are reliant on this axis.

Immunotherapy responses in MSS CRC, which comprise almost 95% of metastatic CRC, are lacking^65^; CAF subpopulations within the TME may be suppressing immune responses in these tumors. Based on our analysis we speculate that targeting ecm-myCAF and TGFβ-myCAF subtypes (responsible for immunotherapy resistance in NSCLC and melanoma) via bispecific antibodies, vaccines, or even cell-based therapies, may enhance current checkpoint blockade strategies^44, 63, 64^. Functional validation and clinical studies will be required to confirm the clinical utility of targeting these CAF populations in CRC.

More importantly, our study’s single-cell resolution enables us to investigate whether tumor cell transcriptomes, and by extension, biological phenotypes, are the primary determinant of CMS classification. Based on our findings, it appears that bulk analysis may have been confounded by varying degrees of tumor microenvironment population enrichment, and that tumor cells within each patient do not segregate into static phenotypes but rather exhibit considerable plasticity. Our single-cell analysis uncovered complex and mixed cellular phenotypes among each cell-specific subpopulation, which projected in a transcriptomic continuum across the CMS groups. These findings were further supported by our scRNA-seq CMS classification analysis that assigned each CRC sample to multiple CMS groups, thereby suggesting CMS heterogeneity within each CRC tumor (Fig. 1d**, Supplementary Table 1**)^16, 66^. These findings may also explain why CMS-defined populations of tumors have not been readily observed in transcriptomic data from independent CRC cohorts^9, 17^. Our results suggest that attempts to divide CRC phenotypes into the current discrete subtypes may undermine optimal patient stratification in the clinical trial setting^9^. The only prospective study to date that utilized the CMS classification (specifically the CMS4 subtype) for patient selection based on dual PD-L1/TGF-ß expression signatures was halted due to futility, suggesting CMS classification may adversely impact clinical trial design^67^.

Previous studies relying on bulk transcriptomics had concerns that stromal content may conceal subtle critical gene signals originating from other key cellular phenotypes within the CRC spectrum, and thus affect CRC classification^8, 9, 50^. Our extensive single-cell analysis allowed for detailed evaluation of all cellular subtypes of CRC simultaneously, thereby uncovering contributions of the different components making up the complex cellular milieu within the CMS classification schema. The present analysis resolved the issue of stromal confounding in CRC and showed that all cellular components contribute to a transcriptomic continuum encompassing all the subtypes that together define the CMS system.

In conclusion, our analysis not only shows tumor-to-tumor variability (as proposed by Ma et al. and other groups), but also demonstrates tumor-TME to tumor-TME as well as TME to TME transcriptional variability at the single-cell resolution level^9, 51^. These data contribute to the conceptual advances in understanding CRC pathogenesis, clinical management, and therapeutic development. We suggest revisiting CMS-like classifications that define CRCs into distinct static subtypes while, in reality CRC tumors exist in a continuum. We caution in using CMS to stratify patients for drug development and clinical trial design. Future studies should concentrate on developing biomarkers (such as CAF’s) and therapeutic agents that can stratify CRC patients beyond traditional classification system boundaries. Ultimately, newer single-cell multiomic technologies will allow us to detect somatic mutations, transcriptomes, proteomes, epigenomes and metabolomes in real-time at the single-cell resolution to better guide more individualized and improved therapies.

## ACKNOWLEDGEMENTS

This study was supported by the startup fund provided to A.M. by the Rush University Medical Center; the OCM grant to A.M by Rush University Cancer Center. Part of A.H.’s time was supported by a Merit Review Award (I01 BX000545) from the Medical Research Service, Department of Veterans Affairs. We would also like to thank Dr. Levi Waldron for sharing code from his publication entitled, “Continuity of transcriptomes among colorectal cancer subtypes based on meta-analysis.” We would also like to thank Dr. Kristian Pietras from providing differential gene expression list of CAFs from his paper titled “Spatially and functionally distinct subclasses of breast cancer-associated fibroblasts revealed by single cell RNA sequencing”. Above all, we want to thank our patients who participated in this study and their families.

## AUTHOR CONTRIBUTIONS

A.M. devised, supervised the study, conducted data analysis, and wrote the manuscript. A.M.K. performed data analyses, wrote the manuscript, and created figures. Z.K. performed data analyses and wrote the manuscript. M.W.G. aided in analysis, wrote the manuscript, and generated figures. A.S. supervised study and wrote manuscript. C.E. and S.S.T. helped with bulk transcriptomic analysis. D.M.H, H.R.G., A.R.B helped with sample collection. All other authors contributed substantially to data interpretation, and manuscript editing. All authors read and approved this manuscript.

## CODE AVAILABILITY

The code generated and utilized in the completion of this publication will be available in a Github repository specific to this project.

## DATA AVAILABILITY

Sequencing data deposition is currently in progress. Ten bulk transcriptomic datasets were accessed from the Gene Expression Omnibus (GEO) database (https://www.ncbi.nlm.nih.gov/geo/).

## METHODS

### Patient and tissue sample collection

Patients with resectable untreated CRC who underwent curative colon resection at Rush University Medical Center (Chicago, IL, USA) were included in this Institutional Review Board (IRB)-approved study. CRC specimens from 16 patients including nine Caucasian, six African American and one Asian patient with corresponding 8 adjacent normal tissue samples were processed immediately after collection at Rush University Medical Center Biorepository and sent for scRNA-seq. Thus, our scRNA-seq atlas represent diverse patient population. The study was conducted in accordance with ethical standards and all patients provided written informed consent.

### Droplet based scRNA-seq-10× library preparation and sequencing

Single-cell RNA sequencing (scRNA-seq) was performed using 10X Genomics Single Cell 5’ Platform. Tumors and normal colon samples were enzymatically dissociated (*Miltenyi*), filtered through a 70-micron cell strainer, pelleted after centrifugation at 300 x*g* and resuspended in DAPI-FACS buffer (PBS, 0.04% BSA). Samples were sorted and viable singlets were gated on the basis of scatter properties and DAPI exclusion. Approximately 3000 cells were pelleted and resuspended in PBS, and cells underwent single cell droplet-based capture on 10X Chromium instruments according to the 10X Genomics Single Cell 5’ Platform protocol. Transcriptome libraries post-fragmentation, end-repair, and A-tailing double-sided size selection, and subsequent adaptor ligation also followed the manufacturer’s protocol. Illumina *NextSeq 550* was used for library sequencing and data were mapped and counted using Cellranger-v3.1.0 (*GRCh38/hg38*).

### scRNA-seq data quality control, gene-expression quantification, dimensionality reduction, and identification of cell clusters

*Cell Ranger* was utilized to process the raw gene expression matrices per samples and all samples from multiple patients were combined in R package (v3.6.3 2020-02-29] -- “*Holding the Windsock*”). Seurat package (v3.2.2) was used in this integrative multimodal analysis^68^. Genes detected in fewer than three cells and cells expressing less than 200 detected genes were filtered out and excluded from analysis. In addition, cells expressing > 25% mitochondria were removed. Cell cycle scoring was performed, (for the S phase and the G2M phase) and the predicted cell cycle phases were calculated. Doublet detection and any higher-order multiplets that were not dissociated during sample preparation were removed via the *DoubletFinder* (v2.0.2) package using default settings^69^. Following quality control one normal colon sample (B-cac13) was discarded due to poor data quality. Finally, 49,859 cells remained and were utilized for downstream analysis.

We adopted the general protocol described in Stuart et al. (2019) to group single cells into different cell subsets^68^. We employed the following steps: clustering the cells within each compartment (including the selection of variable genes for each dataset based on a variance stabilizing transformation [VST]), canonical correlation analysis (CCA) to remove batch effects among the samples, reduction of dimensionality, and projection of cells onto graphs ^70, 71^. Principal component analysis (PCA) was carried out on the scaled data of highly variable genes^72^. The first 30 principal components (PCs) were used to cluster the cells and to perform a subtype analysis by nonlinear dimensionality reduction (t-SNE) and to construct Uniform Manifold Approximation and Projection (UMAP) for cell embeddings^73, 74^. We identified cell clusters under the optimal resolution by a shared nearest neighbor (SNN) modularity optimization-based clustering method. We implemented the *FindClusters* function of the Seurat package, which first calculated *k-nearest neighbors* and constructed the SNN graph. We implemented the original *Louvain algorithm* (algorithm = 1) for modularity optimization. Additionally, we utilized Clustree (v0.4.3) and manual review for identifying the best clustering resolution^19^.

### Major cell type detection and data visualization

To identify all major cell types, we evaluated differentially expressed markers in each identity cell group by comparing them to other clusters using the Seurat *FindAllMarkers* function. We used positively expressed genes with an average expression of >/= 2-fold higher in that subcluster than the average expression in the rest of the other subclusters. We used known marker genes, which have the highest fold expression in that cluster with respect to the other clusters. We also utilized SingleR ((v0.99.10, R Package), which leverage large transcriptomic datasets of well-annotated cell types and manual annotation for cell-type identification^27, 75–77^. Depending on the presence of known marker genes the clusters were grouped as: epithelial cells (*EPCAM, KRT8,* and *KRT18*), fibroblasts (*COL1A1, DCN, COL1A2,* and *C1R*), endothelial cells (CD31+), myeloid cells (*LYZ, MARCO, CD68,* and *FCGR3A*), CD4 T cells (*CD4*), CD8 T cells (*CD8A* and *CD8B*), and B cells (*MZB1*), ^27, 39, 43, 75, 78–81^. The cells were eventually assembled into DGE matrices within each compartment, containing all six cell types.

### Major-cell type subclustering and data visualization

Each major cell type, including epithelial cells, endothelial cells, T cells, B cells, myeloid cells, and fibroblasts was reclustered and reanalyzed to study each compartment at a higher resolution to detect granular cellular heterogeneity in CRC. Clustree (v0.4.3) and manual review were utilized for optimal cluster detection. For cell annotation of each cell type, we utilized published literature gene expression signatures and manual review of differential genes among clusters. Additionally, we again utilized SingleR (v0.99.10, R Package) for unbiased cell annotation. Interestingly, reclustering of major compartments individually also detected clusters expressing hybrid markers as well as cell clusters expressing markers from distinct lineages (such as T cell clusters expressing B cells); these were removed and excluded for further analysis. We utilized UMAP for visualization purposes. For validation, we analyzed 65,362 cells from 23 patients and applied the similar quality control metrics as outlined above. In addition, we also applied *vars.to.regress* function to remove low quality cells in an unbiased manner. We retained 31,383 high-quality single cells for further analysis^25^. These high-quality cells were analyzed utilizing the same pipelines and parameters as that for our primary cohort (Supplementary Figs. 4-5 and 11-12).

The InferCNV (v1.2.1) package was used with default paramets to identify the evidence for somatic large-scale chromosomal copy number alteration in epithelial cells (*EPCAM+, KRT8+, KRT18+*)^82^. Normal epithelial cells were used as the control group.

### Trajectory analysis

We used Monocle v.2 (v2.14.0), a reverse graph embedding method to reconstruct single-cell trajectories in tumor and normal epithelium^83^. In brief, we used UMI count matrices and the *negbinomial.size*() parameter to create a *CellDataSet* object in the default setting. We grouped projected cells on UMAP in default settings for visualization of monocle results. We defined the cumulative duration of the trajectory to show the average amount of transcriptional transition that a cell undergoes as it passes from the starting state to the end state. The cells were also ordered in pseudotime to explain the transition of cells from one state to another.

### Pathway-Gene set variation analysis (GSVA)

Pathway analysis was performed on the 50 hallmark gene sets downloaded from *Molecular Signatures Database (v7.2)*. We used GSVA (v1.34.0), a non-parametric, unsupervised method to estimate the gene set variations and evaluation of pathway enrichment, and pathway scores were calculated for each cell using standard settings ^29, 33^.

### DNA and bulk RNA library construction

DNA and bulk RNA sequencing was performed as previously described^84^. One hundred nanograms of DNA from each tumor was mechanically sheared to an average size of 200 bp. Using the *KAPA Hyper Prep Pack*, DNA libraries were packed, hybridized into the *xT probe* package, and amplified with the *KAPA HiFi HotStart ReadyMix*. For uniformity, each sample needed to have 95% of all targeted base pairs sequenced to a minimum depth of 300x. One hundred nanograms of RNA per tumor sample was heat fragmented to a mean size of 200 base pairs in the presence of magnesium. Using random primers, the RNA was used for first-strand cDNA synthesis, followed by second-strand synthesis and A-tailing, adapter ligation, bead-based cleanup, and amplification of the library. After library planning, the *IDT xGEN Exome Test Panel* was hybridized with samples. Streptavidin-coated beads and target recovery were carried out, accompanied by amplification using the *KAPA HiFi* library amplification package. The RNA libraries were sequenced on an *Illumina HiSeq 4000* using patterned flow cell technology to achieve at least 50 million reads.

### Detection of somatic variation on DNA sequencing data

The tumor and normal FASTQ files were paired. For quality management measurement, FASTQ files were evaluated using FASTQC and matched with Novoalign (Novocraft, Inc.)^84, 85^. SAM files were generated and converted to BAM files. The BAM files were sorted, and duplicates were marked. Single nucleotide variations (SNVs) were called after alignment and sorting. For discovery of copy number alterations, the de-duplicated BAM files and the VCF generated from the variant calling pipeline were processed to computate read depth and variance of heterozygous germline SNVs between the tumor sample and normal sample. Binary circular segmentation was introduced and segments with strongly differential log_2_ ratios between the tumor and its comparator were chosen. From a combination of differential coverage in segmented regions and estimation of stromal admixture provided by analysis of heterozygous germline SNVs, an estimated integer copy number was determined

### Microsatellite instability status

Probes for 43 microsatellite regions were developed using *Tempus xT* assay^84^. Tumors were categorized into three groups by the MSI classification algorithm as described by Tempus: microsatellite instability-high (MSI-H), microsatellite stable (MSS) or microsatellite equivocal (MSE). MSI screening for paired tumor-normal patients used reads mapped to the microsatellite loci with at least 5 bps flanking the microsatellite. The sample was graded as MSI-H if there was a >70% chance of MSI-H classification. If the likelihood of MSI-H status was 30-70%, the test findings were too ambiguous to interpret and those samples were listed as MSE. If there was a <30% chance of MSI-H status, the sample was called MSS. Additionally, IHC results were used to classify tumors into MSS or MSI molecular subtypes. Both of these modalities were concordant and produced the same results.

### Bulk transcriptomics analysis

We downloaded and pooled eight colorectal gene expression datasets (GSE13067^86^, GSE13294^86^, GSE14333^48^, GSE17536^49^, GSE20916^87^, GSE33113^88^, GSE35896^89^, and GSE39582^11^), a pancreatic cancer dataset (GSE62165^90^) and a non-small cell lung cancer dataset (GSE33532^91^) to validate our findings from the single cell compartments by deconvoluting the bulk gene expression profiles into pseudo single-cell resolutions. We used Affy (v1.64.0) for the data analysis and for exploration of Affymetrix oligonucleotide array probe level data^92^. Batch correction was carried out using the removeBatchEffect (v3.42.2) function of the LIMMA program and CMScaller for the CMS classification (see below)^93^. Three datasets (GSE17536^49^, GSE33113^88^, and GSE39582^11^) were utilized for clinical outcome analysis^93, 94^.

### Correlation patterns in bulk gene expressions for CAF compartments

To identify the top correlated CAF-marker genes within the combined eight CRC datasets, three bulk CRC gene expression sets individually, pancreas cancer and lung cancer datasets. We first transformed the bulk gene expression sets with log_2_ transformation. Next, marker genes with an average log_2_ FC>/=0.5 and p<0.05 obtained from the single cell data of CAF-S1 and CAF-S4 compartments were separately intersected with the bulk gene expression sets. Genes with an average Spearman correlation score greater than 0.8 were kept as the CAF signatures within the bulk gene expression. Heatmaps illustrating the correlation patterns within and between the CAF compartments were prepared with the heatmap.2 function from ggplot package (v3.1.1) utilizing the Pearson correlation coefficient. Heatmaps illustrating the correlation patterns within and between the CAF compartments were prepared using the ggplot package (v3.1.1) utilizing the Pearson correlation coefficient^95^.

The Cox proportional hazard regression model was used to examine the significance of 20 cell types from scRNA-seq in bulk expression data. Each cell type’s marker genes with an average logFC>1 and adjusted P<0.05 were intersected with the bulk expression datasets separately. We only kept the marker genes with a high correlation with each other in bulk, which provides an average correlation score of > 0.8. The average bulk expression of each cell type’s remaining marker genes was calculated and used in the hazard regression model as the representative of this cell type. For analysis of relationships with patient outcome, univariate models were calculated using Cox proportional hazard regression (coxph function from survival R package)^96^.

### Deconvoluting public bulk gene expression profiles into pseudo single-cell expressions

We used CIBERSORTx v1 to estimate composition of various cell populations in GSE39582^11^, GSE14333^48^, and GSE17536^49^ datasets^47^. Signature gene matrices were created using the expression profiles of 49,859 cells as the reference single cell profile. We ran the ‘hires’ module with default parameters except for the ‘rmbatchBmode,’ and the bulk-mode batch correction argument was set to true. After the deconvolution process, we normalized the gene expressions according to the cell fractions in each sample and calculated each gene’s Z-transformed expression values. The average normalized expression of each cell type across all samples was plotted with the heatmap.3 R function of the GMD package (v0.3.3)^97^. A signature matrix highlighting marker genes of the different cell types was prepared with a heatmap.2 R function of ggplot (v3.1.1).

### Consensus molecular subtyping of colorectal cancer (CMS Classification)

We used R package CMScaller(v0.9.2), a nearest template prediction (NTP) algorithm, for the classification of gene expression datasets^94^. We set the permutation number to 1000 to predict the CMS classes of the samples in the GEO datasets with a p-value < 0.05. We ran CMScaller with default parameters.

### Continuous subtype discovery using scRNA-seq analysis

Bulk mRNA expression profiles of the GEO dataset (GSE39582^11^), composed of 585 samples in total, were deconvoluted into the pseudo single-cell expression profiles via CIBERSORTx utilizing the expression data consisting of 20 different well known cell types from our scRNA-seq dataset^47^. We transformed the deconvoluted expression matrix with log2 transformation. The principal components cluster subtype scores (PCSSs) of the CMS among the samples, were determined separately for each cell type using an algorithm published by Ma et al^51^. To obtain the PCSSs, the average loading vectors were used. The results obtained for 20 cell types were projected on the first two PCSSs (PCSS1 and PCSS2) as they were validated by Ma et al. in their analysis using 18 datasets. We also analyzed two datasets (GSE14333^48^, GSE17536^49^) to independently confirm reproducibility of continuous scores.

### Statistics and reproducibility

All statistical analyses and graphs were created in R (v3.6.3) and using a Python-based computational analysis tool. Schematic representations were made using the Inkscape (https://inkscape.org/) software. Dim plots, bar plots and box plots were generated using the dittoSeq (v1.1.7) package with default parameters^98^. Violin plots were generated using the patchwork (v1.1.0) package and ggplot2 (v3.3.2) package in R with default parameters. Heatmaps were generated using Morpheus.R with default parameters^99, 100^. ANOVA and pair-wised t-tests for the CMS classes across the deconvoluted expression profiles were performed in R using the ggpubr R (v0.4.0) package^101^. The Box and Whisker plots were generated using the boxplot function of the R base package at default parameters. The mean of the log_2_ transformed deconvoluted expression value of the samples in each CMS group was demonstrated with a horizontal straight line within each box. The length of a boxplot corresponds to the interquartile range (IQR), which is defined as the range between the first and third quartiles (Q1 and Q3), whereas the whiskers are the upper and lower extreme values of the data (either data’s extremum values, or the Q3+1.5*IQR and Q1-1.5*IQR values, whichever was less extreme). To test for differential score based on CIBERSORTx cell abundances between different molecular subtypes pairwise t-test with Benjamini-Hochberg correction was used.

### Survival analysis

Survival curves were obtained according to the Kaplan-Meier method survfit (v3.2-7), and differences between survival distributions were assessed by Log-rank test. The patients were divided into two groups (high/poor and low/good risk) according to their median expression values (survminer (v0.4.8)). The surv_cutpoint function uses the maximally selected rank statistics and implements standard methods for the approximation of the null distribution of maximally selected rank statistics (maxstat (v0.7-25)).

The proportional hazard assumption was tested to examine the fit of the model for survival of the samples in three GEO datasets (GSE17536^49^, GSE33113^88^, and GSE39582^11^) with respect to the deconvoluted bulk mRNA expressions. For analysis of the relationships with patient outcome, multivariate models were calculated using the Cox proportional hazard regression (coxph survival R package)^96^.

## Supplementary Materials

**Supplementary figure 1.**
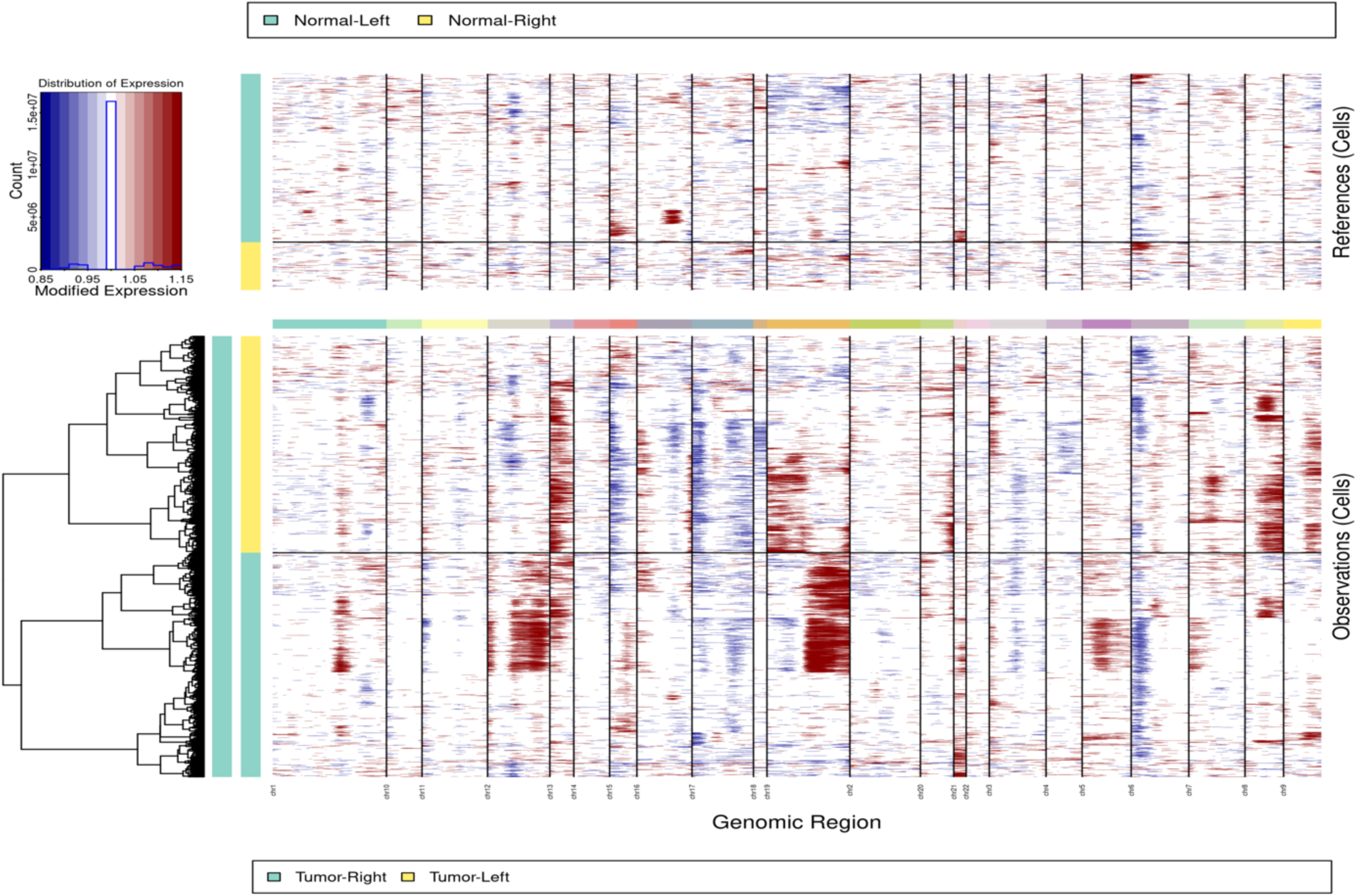
Analysis of copy number variation amongst epithelial cells. CNV analysis was conducted on the epithelial cell compartment. Increase CNV is seen in tumor derived epithelial cells (observation). Non-malignant derived epithelial cells were used as control (references).

**Supplementary figure 2.**
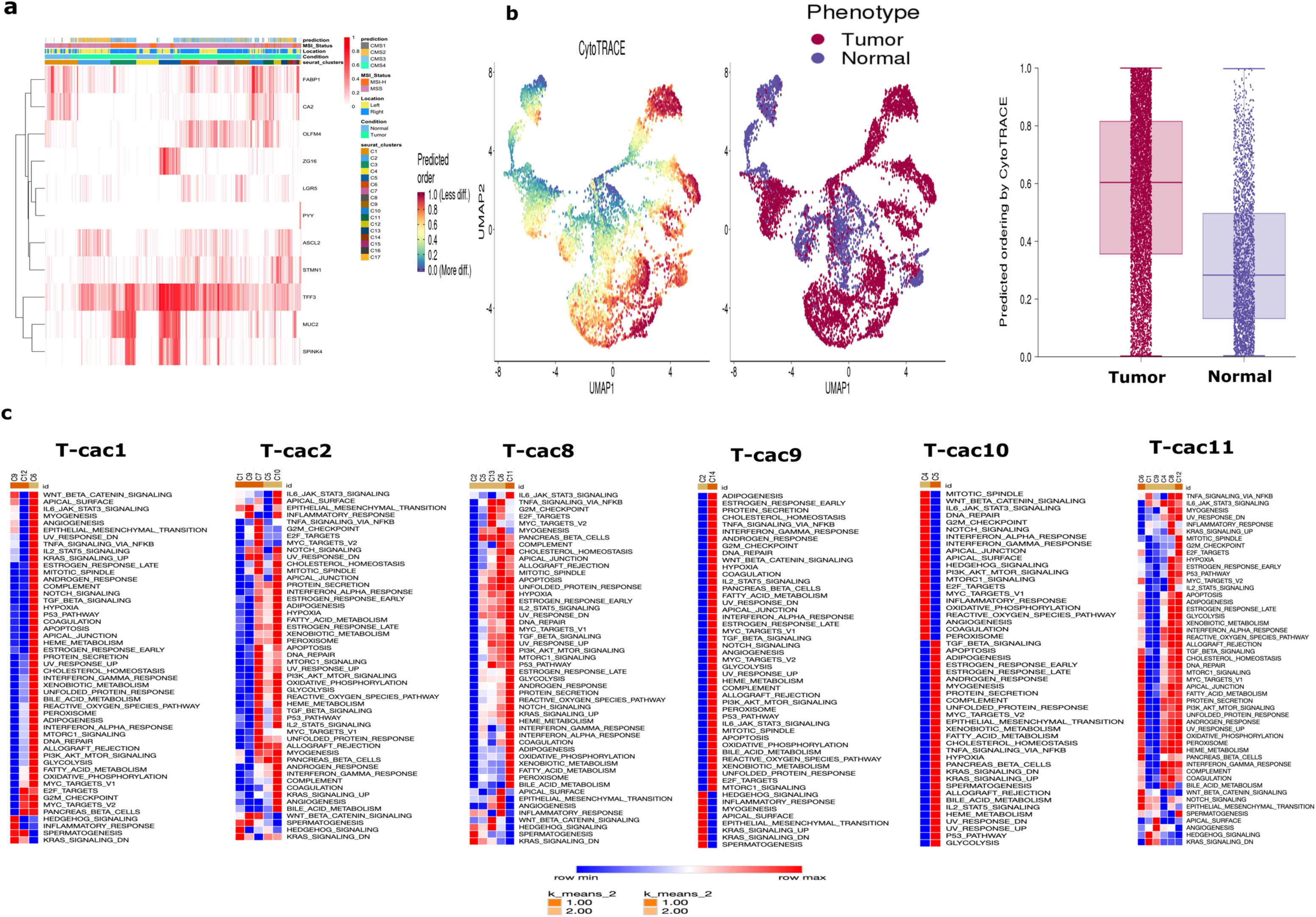
**a,** Heatmap of marker gene expression for the epithelial and tumor cells. **b,** UMAP visualization of computational analysis of differentiation status using CytoTRACE (see methods). **c,** Heatmap representation of Hallmark pathway analysis of epithelial phenotypes within six different tumor samples demonstrating subclonal intratumoral heterogeneity.

**Supplementary figure 3.**
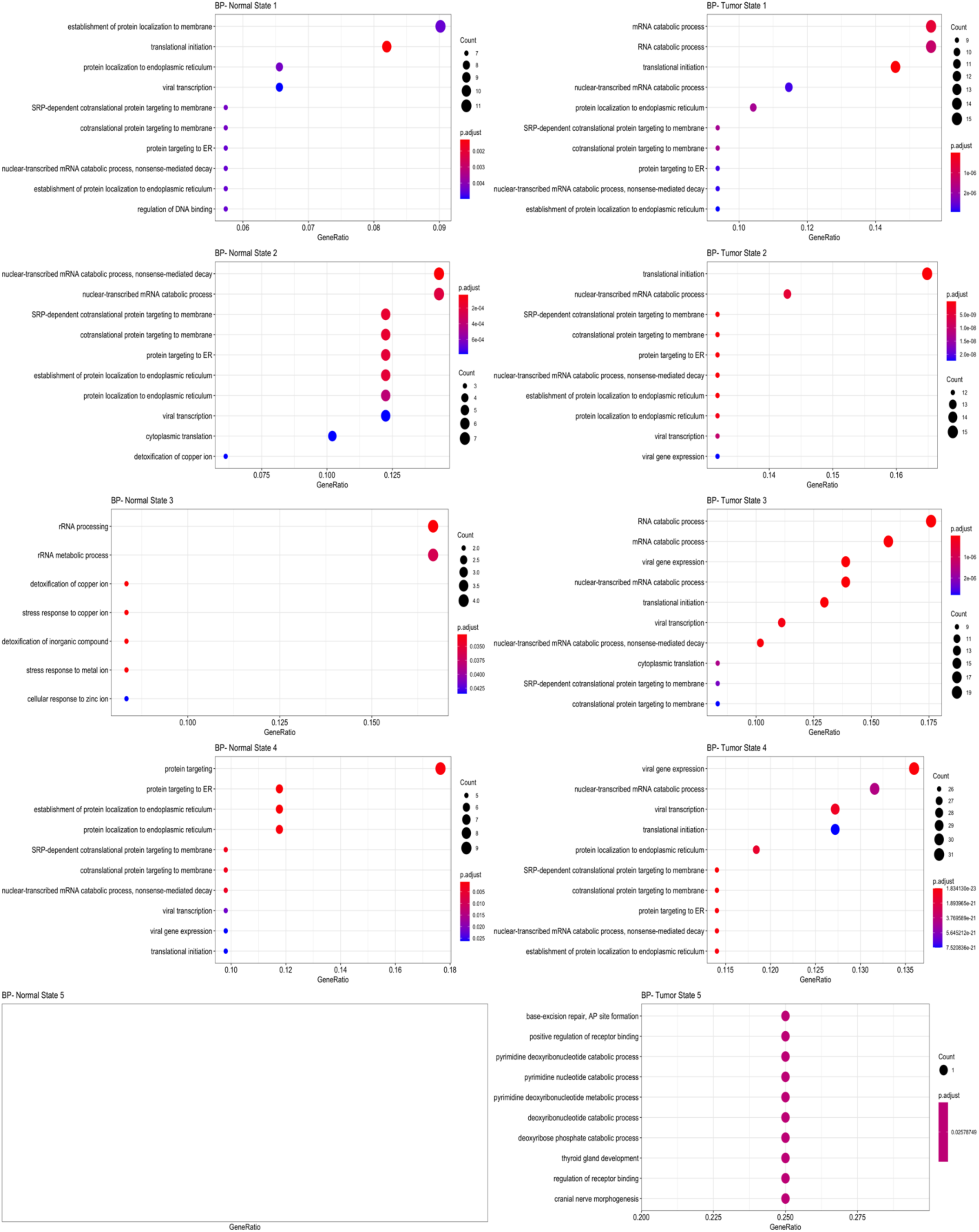
Pathway analysis (GO ontology) of gene-expressions specific to each cell-state in trajectory analysis stratified by malignant and non-malignant sample of origin.

**Supplementary figure 4.**
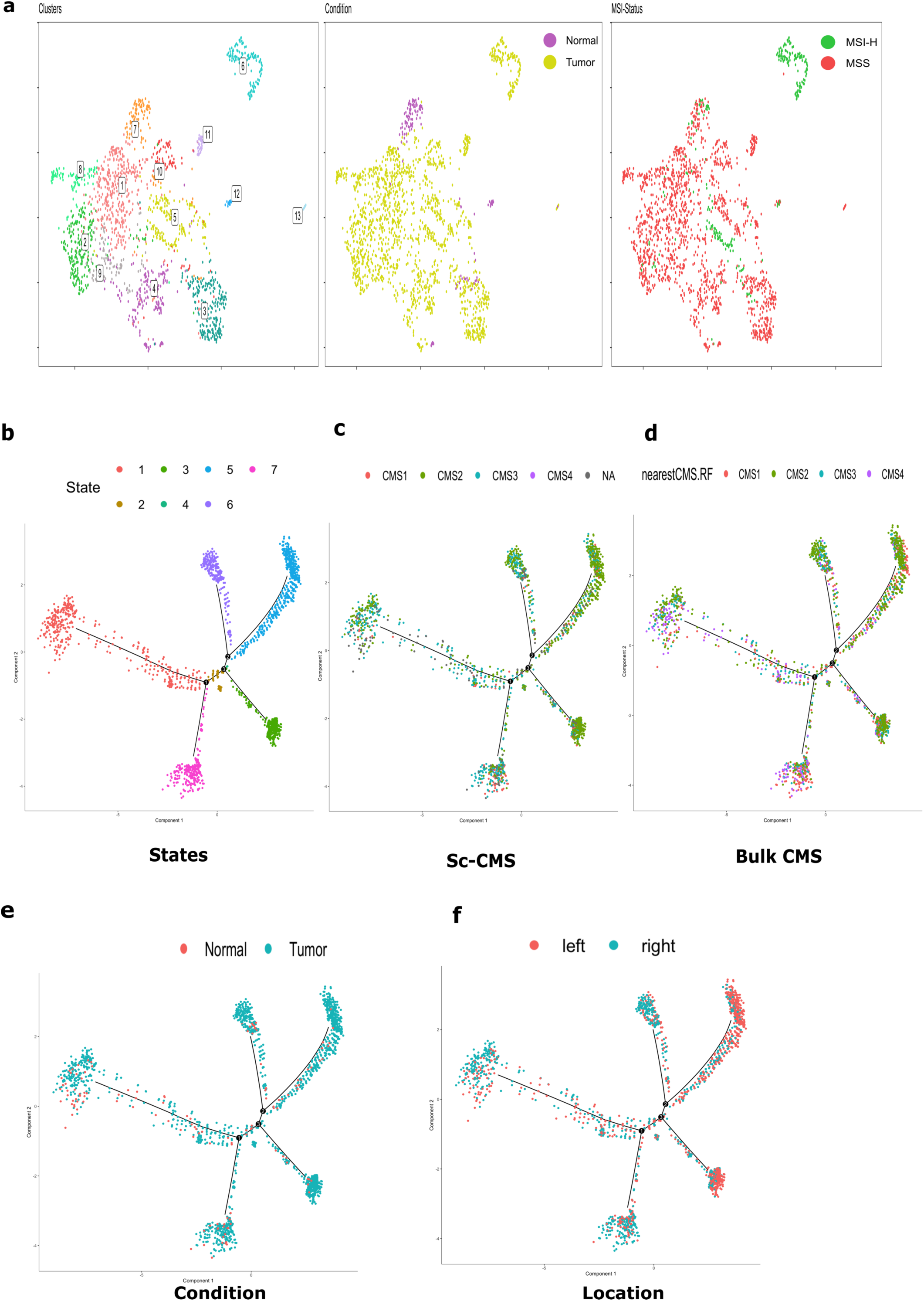
Epithelial clustering and differentiation trajectories from a validating cohort (Lee et al. data). **a,** UMAP clustering of epithelial cells colored by cluster, sample of origin tumor vs. normal tissue status and MSI status. **b-f,** Differentiation trajectories of epithelial cells colored by differentiation state, CMS status, tumor status and colonic location.

**Supplementary figure 5.**
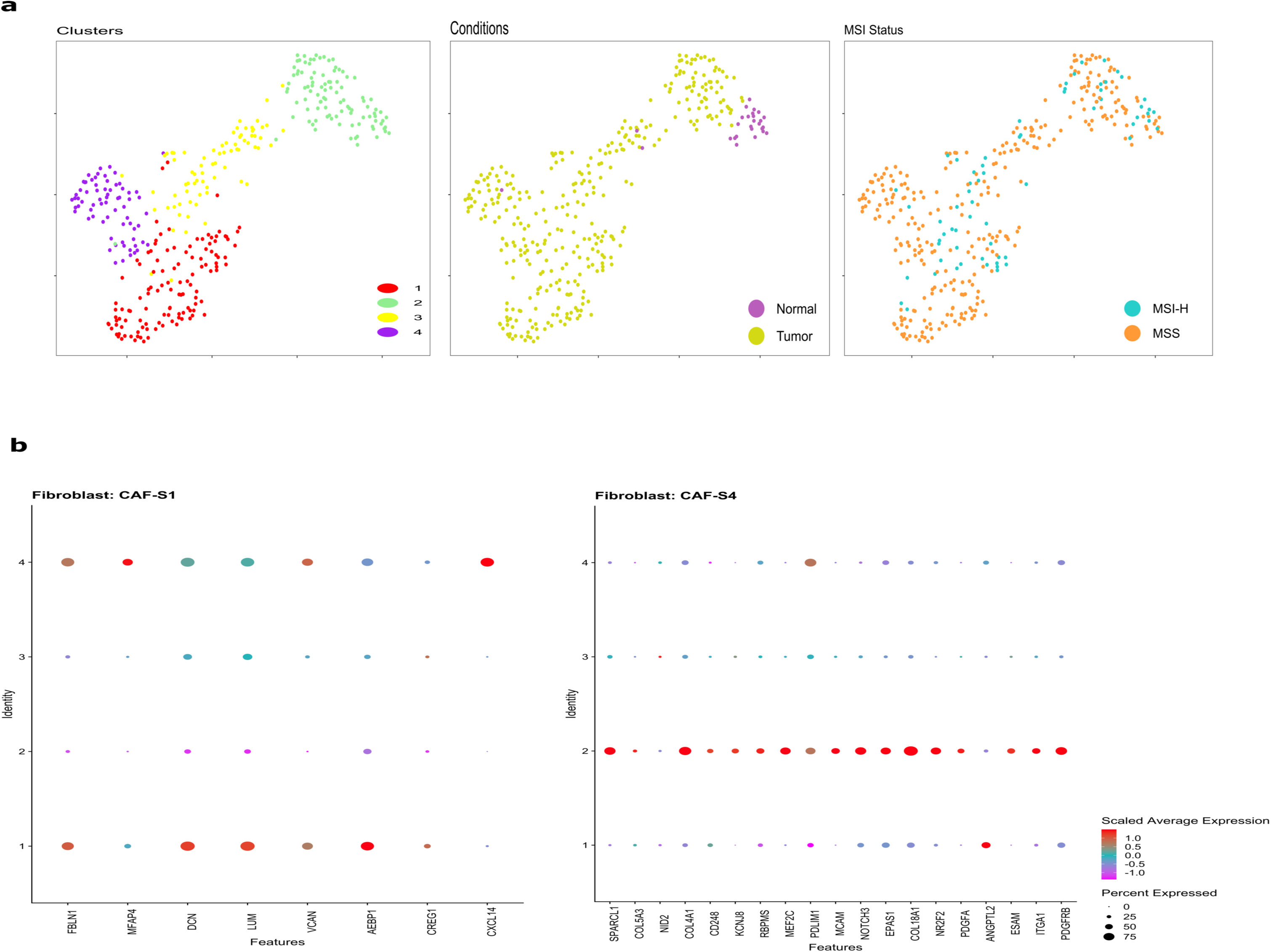
Characterization of fibroblasts and their transcriptomic expression patterns (Lee et al. data). **a,** Fibroblasts colored by distinct groups, tumor vs normal sample, and sample specimen. **b,** Dot plot demonstrating variable expression patterns of subtypes of CAF-S1 and CAF-S4 confirming their relevance in colorectal cancer.

**Supplementary Figure 6.**
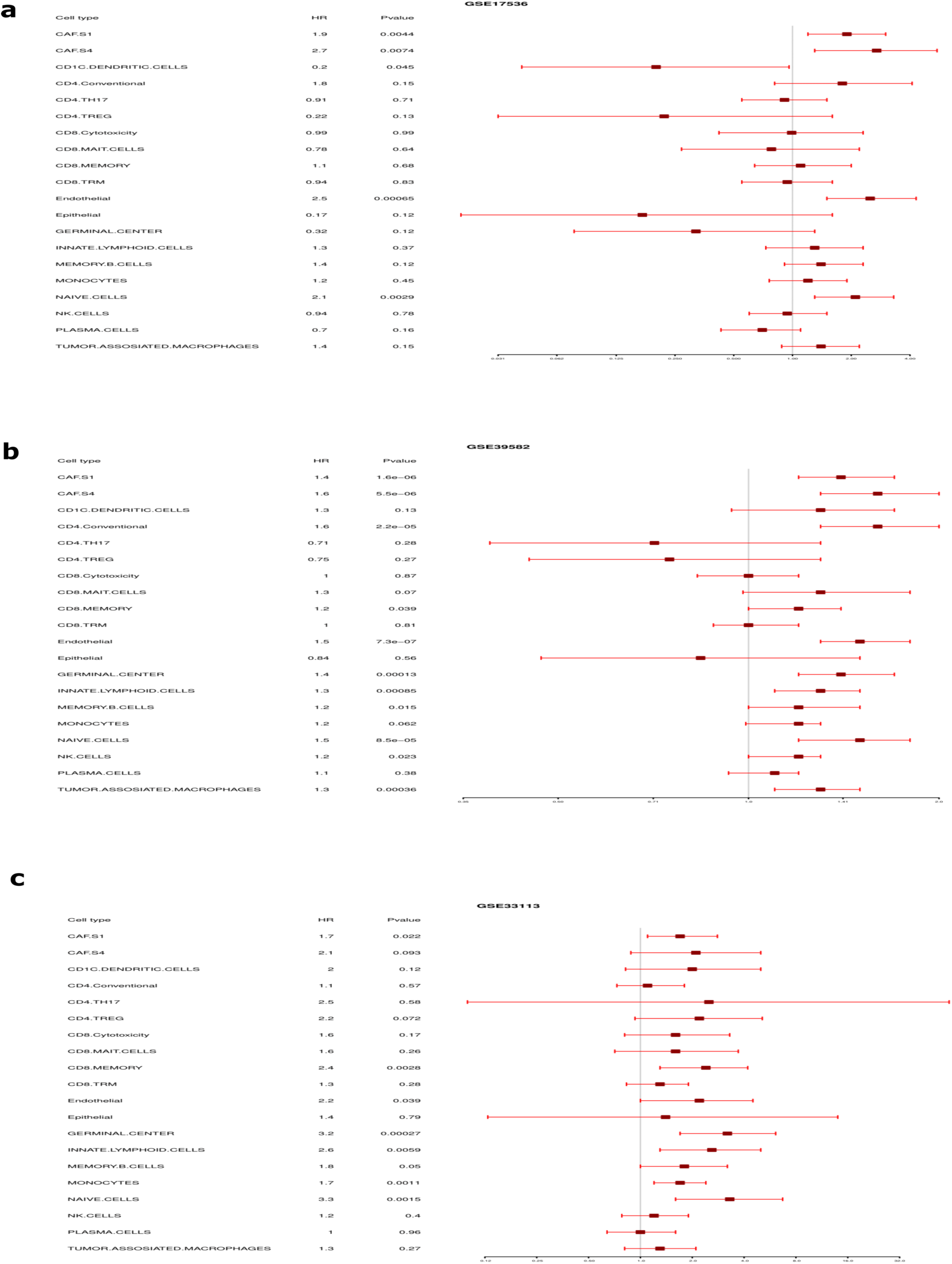
Association between relative cell abundance and patient survival from microarray-based datasets. a. GSE17536 (n=177). b GSE39582 (n=585). c. GSE33113 (n=96). Note that CAF-S4 is not significant in GSE33113 (p=0.093).

**Supplementary Figure 7.**
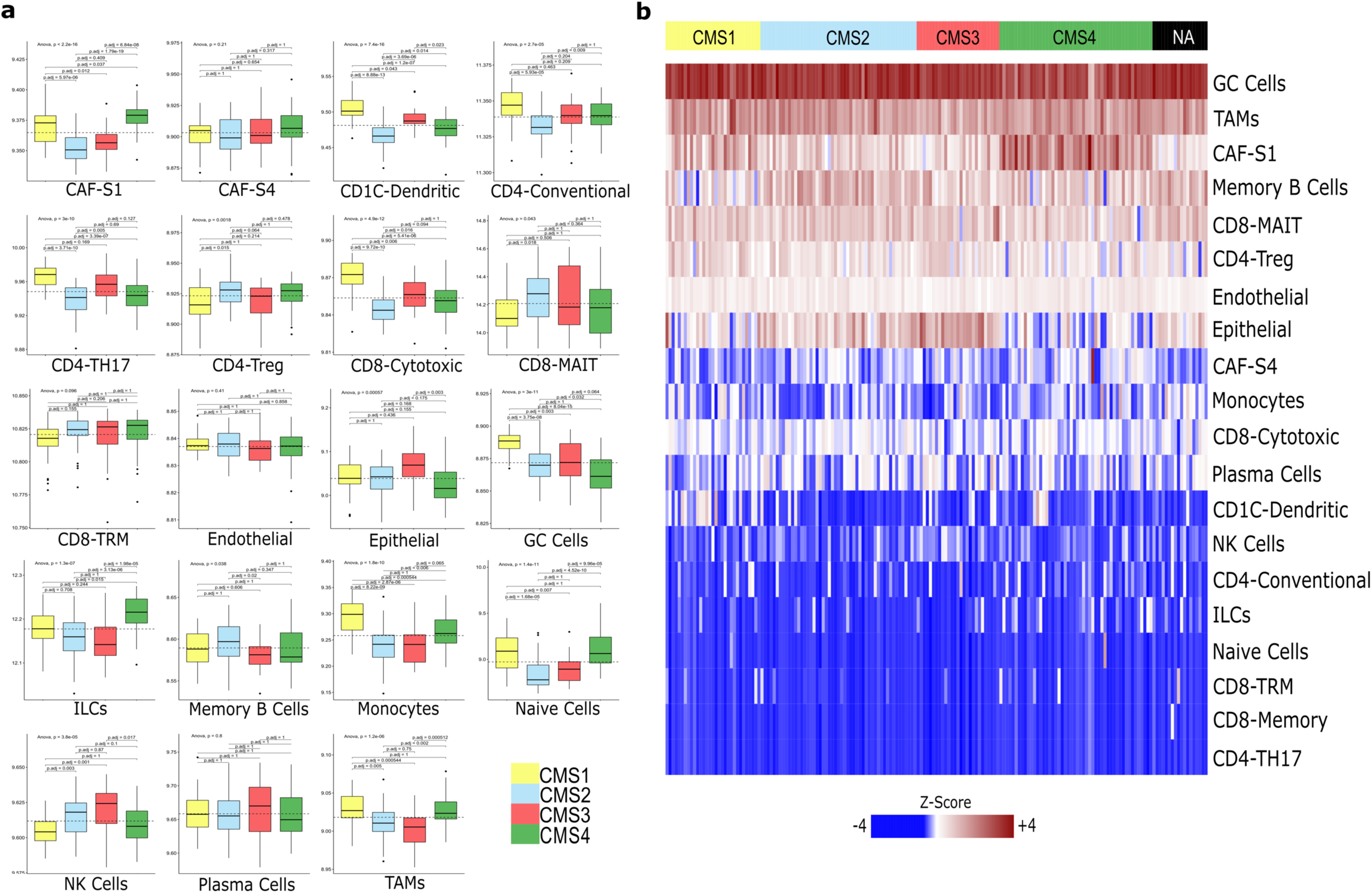
Cell abundance prediction for each sample and projected on bulk CMS on GSE17536 (n= 177). **a,** Boxplots show the distribution of cell types within tumors with varying CMS status. The whiskers depict the 1.5 x IQR. The p-values for pairwise t-tests comparisons (with Benjamani-Hochberg correction) and ANOVA tests of cell abundance across CMS are shown in the figure. **b,** Deconvolution heatmap of different cell types by average expression using CIBERSORTx demonstrating cell type distribution within each CMS category. ILCs = Innate lymphoid cells, GC Cells = Germinal Center B Cells, NK Cells = Natural Killer Cells, TAMs = Tumor Associated Macrophages.

**Supplementary Figure 8.**
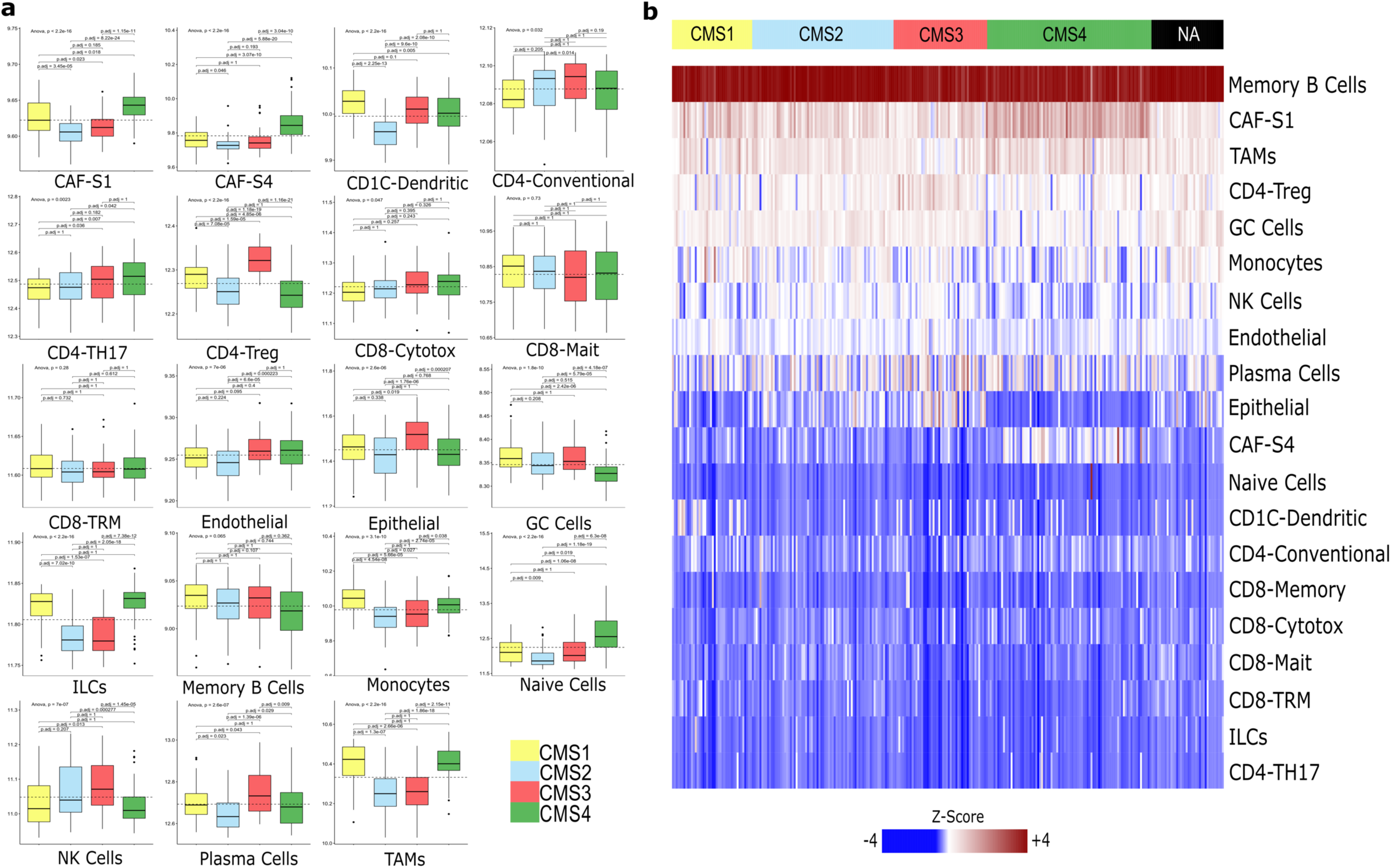
Cell abundance prediction for each sample and projected on bulk CMS on GSE14333 (n= 290). **a,** Boxplots show the distribution of cell types within tumors with varying CMS status. The whiskers depict the 1.5 x IQR. The p-values for pairwise t-tests comparisons (with Benjamani-Hochberg correction) and ANOVA tests of cell abundance across CMS are shown in the figure. **b,** Deconvolution heatmap of different cell types by average expression using CIBERSORTx demonstrating cell type distribution within each CMS category. ILCs = Innate lymphoid cells, GC Cells = Germinal Center B Cells, NK Cells = Natural Killer Cells, TAMs = Tumor Associated Macrophages.

**Supplementary Figure 9.**
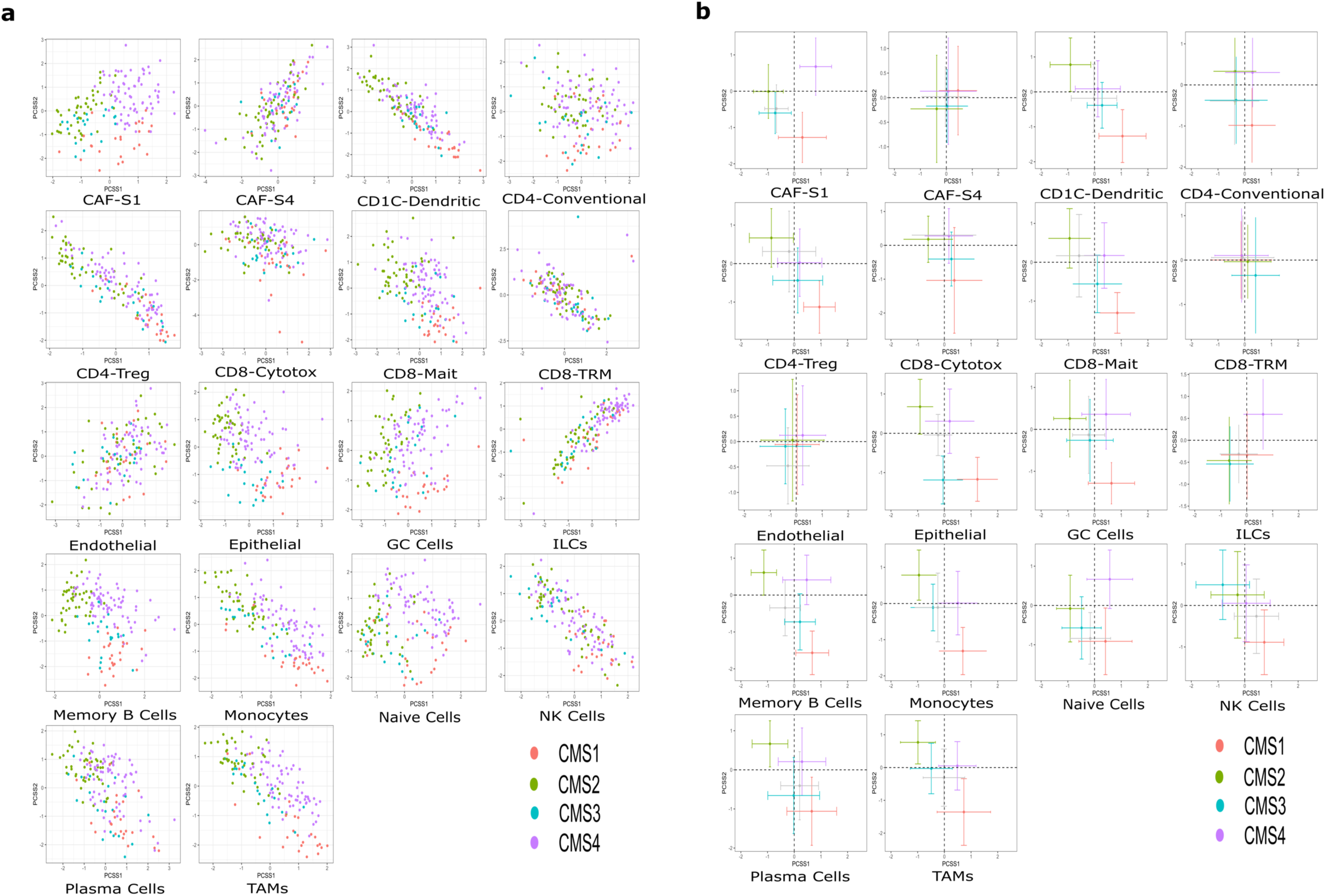
Continuous scores for CRC dataset GSE17536 (n= 177). **a,** Principal component analysis plot showing PCSS1 and PCSS2 continuous scores reported by CMS classification across cell types show minimal separation in the top 2 principal components. **b,** All cell types projected on four quadrants representing CMS1-4 using PCSS1 and PCSS2 scores. Note that the cell types largely form a continuum along CMS status and are not clustered in discrete quadrants separate from one another. Cells and markers are colored by bulk CMS status accordingly to the tumor sample of origin. ILCs = Innate lymphoid cells, GC Cells = Germinal Center B Cells, NK Cells = Natural Killer Cells, TAMs = Tumor Associated Macrophages.

**Supplementary Figure 10.**
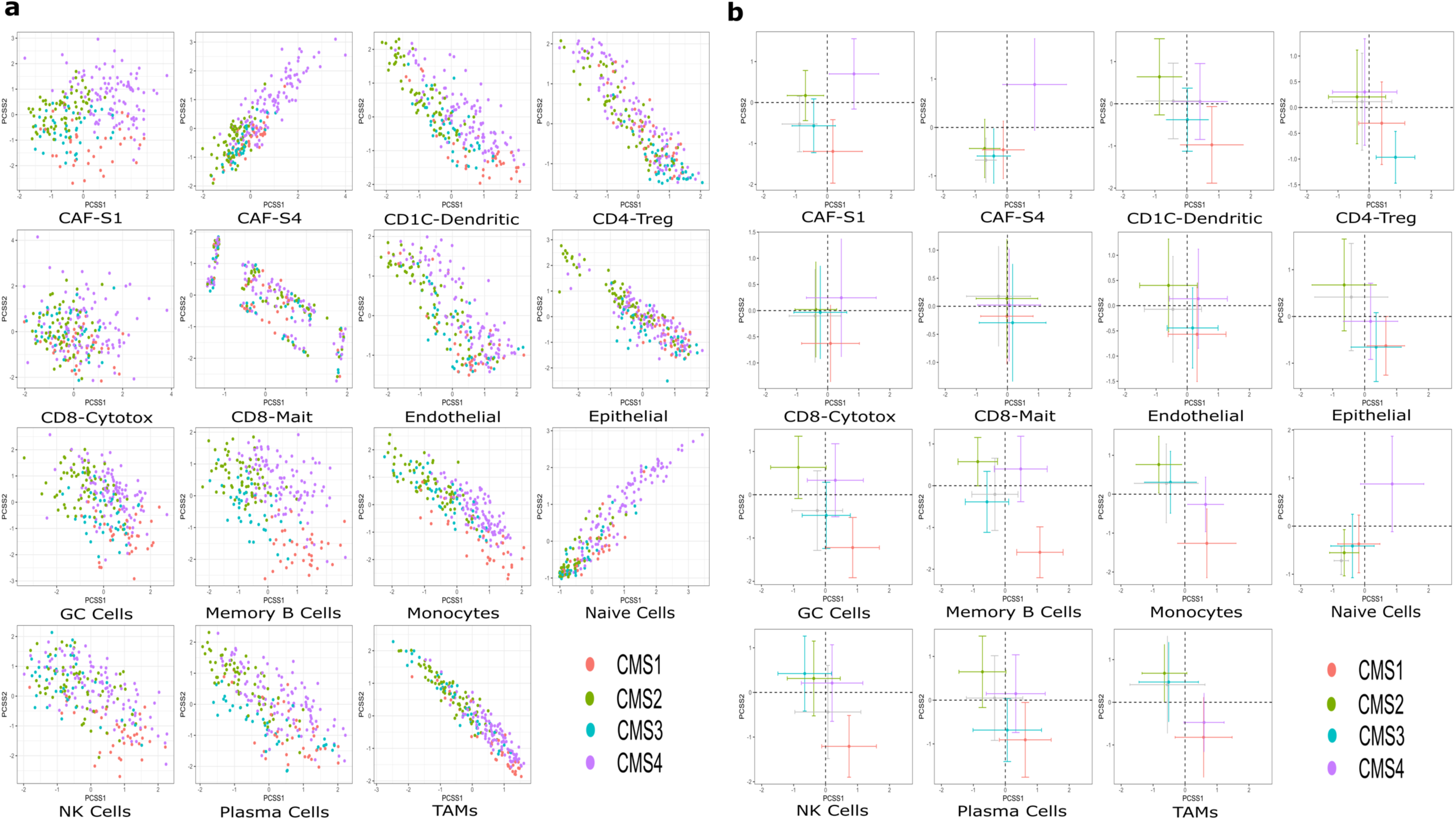
Continuous scores for CRC dataset GSE14333 (n= 290). **a,** Principal component analysis plot showing PCSS1 and PCSS2 continuous scores reported by CMS classification across cell types show minimal separation in the top 2 principal components. **b,** All cell types projected on four quadrants representing CMS1-4 using PCSS1 and PCSS2 scores. Note that the cell types largely form a continuum along CMS status and are not clustered in discrete quadrants separate from one another. Cells and markers are colored by bulk CMS status accordingly to the tumor sample of origin. ILCs = Innate lymphoid cells, GC Cells = Germinal Center B Cells, NK Cells = Natural Killer Cells, TAMs = Tumor Associated Macrophages.

**Supplementary figure 9.**
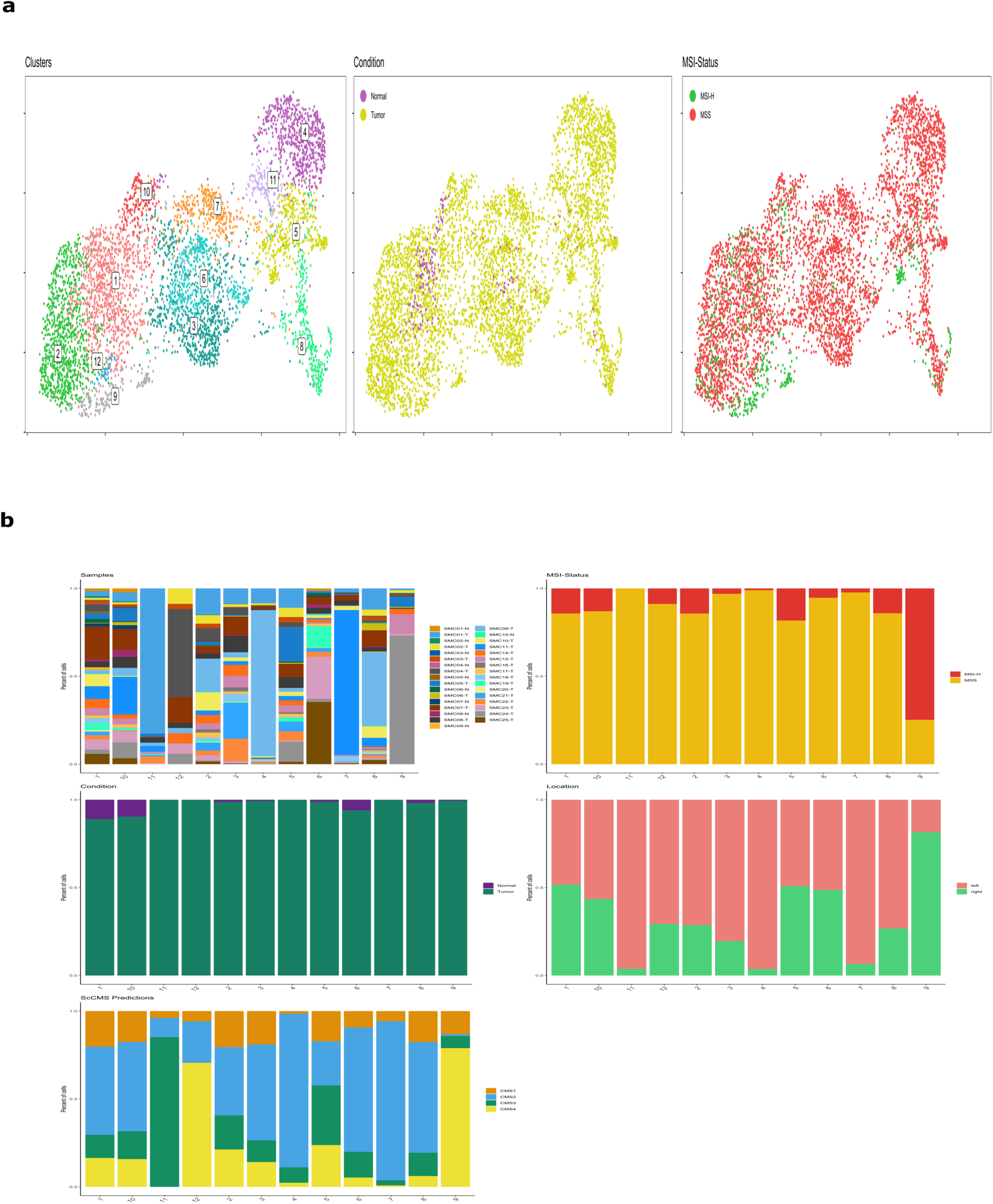
T/NK cell data from Lee et al. **a**, Reclustering of T/NK cells and coloring by clusters, tumor and normal status, MSI status and samples. **b**, Bar chart representation of cells colorized by our samples of origin, MSI status, tumor status, colonic location, CMS status.

**Supplementary figure 10.**
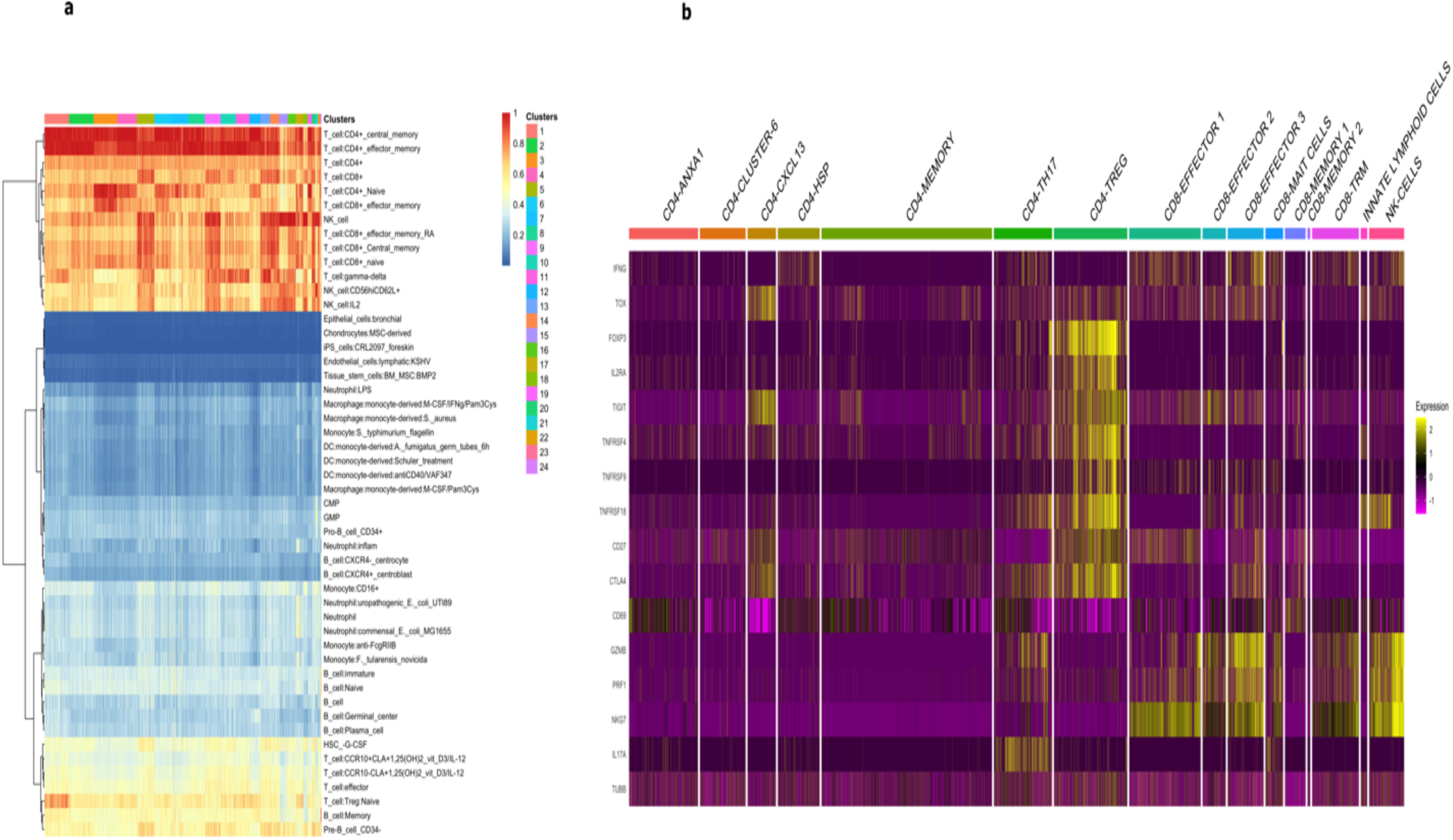
T cell expression analyses across T/NK cells. **a,** SingleR heatmap cell type identification within each cluster. Note that genes associated with T cells show increased expressions, confirming the quality of the data. Note doublets were removed from the further analysis. **b,** T/NK cells gene specific expression to identify T cell heterogeneity using published literature (see methods).

**Extended Data Figure 1.**
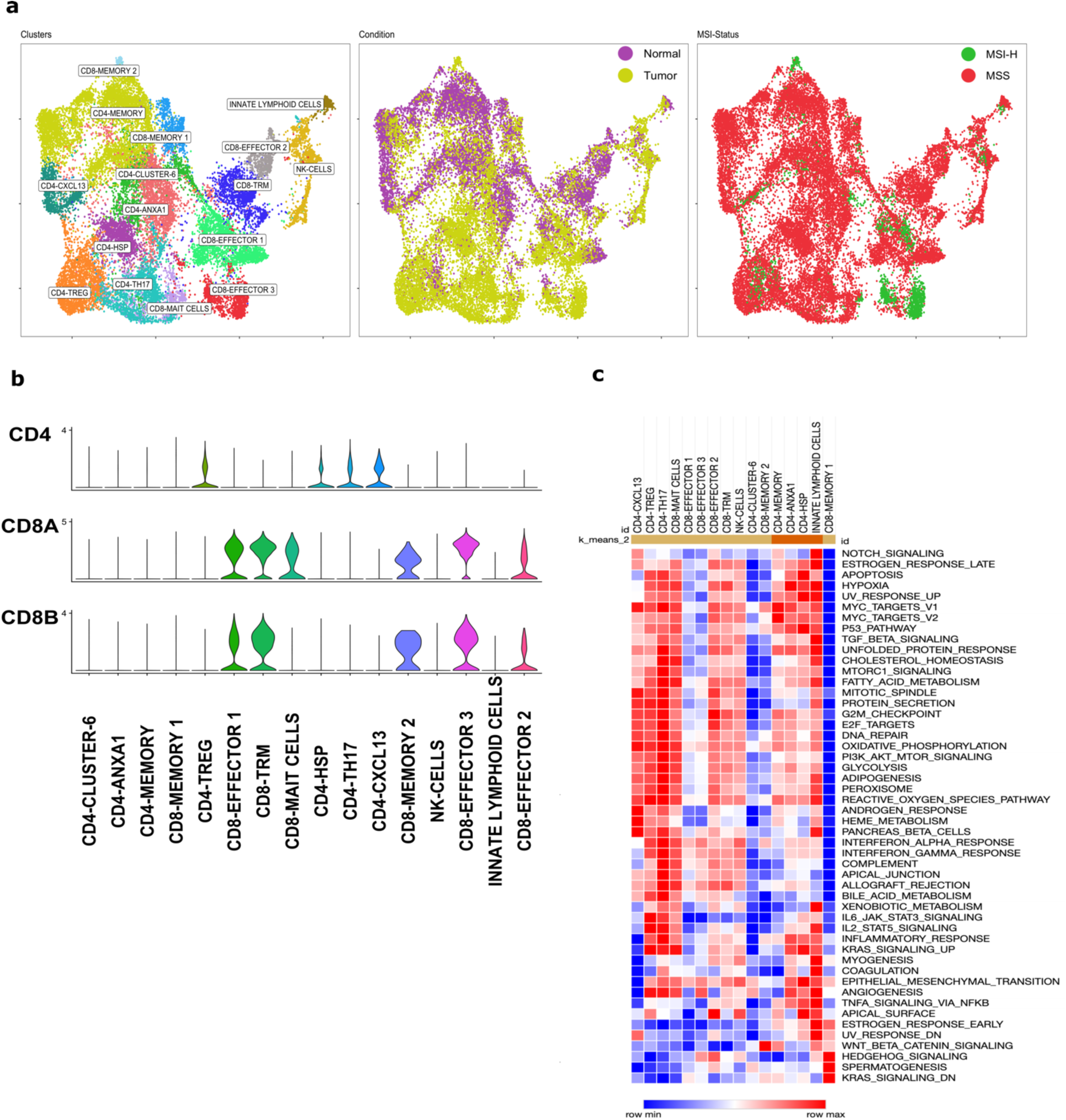
T cell identification and characterization. **a**, UMAP reclustering of T cells colored by cell phenotype, tissue malignancy status, and sample of origin. **b**, Violin plots showing the differential expression of T cell-specific marker genes between CD4 and CD8 phenotypes. **c**, Heatmap of the Hallmark pathway analysis for the T cell compartment. scCMS, single-cell consensus molecular subtyping; bulk CMS, consensus molecular subtyping on Bulk RNA-seq data. **Description:** We reclustered and analyzed 22,525 cells from both tumors and adjacent, normal tissue samples and identified 11 CD4+ T cell and 12 CD8+ T cell clusters. We used known canonical markers and published expression signatures to identify T cell states for further analysis (see methods). We identified conventional CD4+ T cells, CD4+ Tregs, CD8+ (naive/memory, cytotoxic, resident memory, and MAIT cells), NK cells, and innate lymphoid cells (ILC). Among the conventional CD4+ T cells, we identified the central memory/naive like state (CCR7+, SELL+, and TCF7+) enriched in non-malignant samples. In contrast, Th17 cells expressing IL-17, known as critical anti-tumor effectors, were enriched in tumor samples. CD4+ Tregs (FOXP3+) expressing immune checkpoint markers and costimulatory molecules were among the most abundant T cells in the colorectal TME compared to non-malignant tissue. Among the CD8+ T cell states, CD8+ cytotoxic cells were distributed across three clusters that we labeled CD8+ effector 1, CD8+ effector 2, and CD8+ effector 3. CD8+ effector clusters expressed cytotoxicity genes and chemokines as previously described in other tumor types. CD8 effector3 was predominantly enriched in MSI-H CRC patients and represented 77% of the total CD8+ effector 3 population among the 2 MSI-H CRC samples. This cluster expressed ITGAE, LAYN, CXCL13, and T cell exhaustion markers (LAG3, HAVCR2, and CD96), possibly explaining this CD8+ cell state’s role in the response to immune checkpoint inhibitors in MSI-H colorectal tumors. Gene-set enrichment of CD8+ effectors further confirmed their distinct states. CD8+ effector 2 was a proliferative cluster with MYC activity, NOTCH activation, and EF2 targets. CD8+ effector(s), CD8+ MAIT cells, and NK cells were enriched in tumors, whereas Tissue-resident memory (Trm) cells were depleted in tumor tissue. Trm induction was recently seen to enhance cancer vaccine efficacy in other tumors, suggesting a possible therapeutic target in CRC.

**Extended Data Figure 2.**
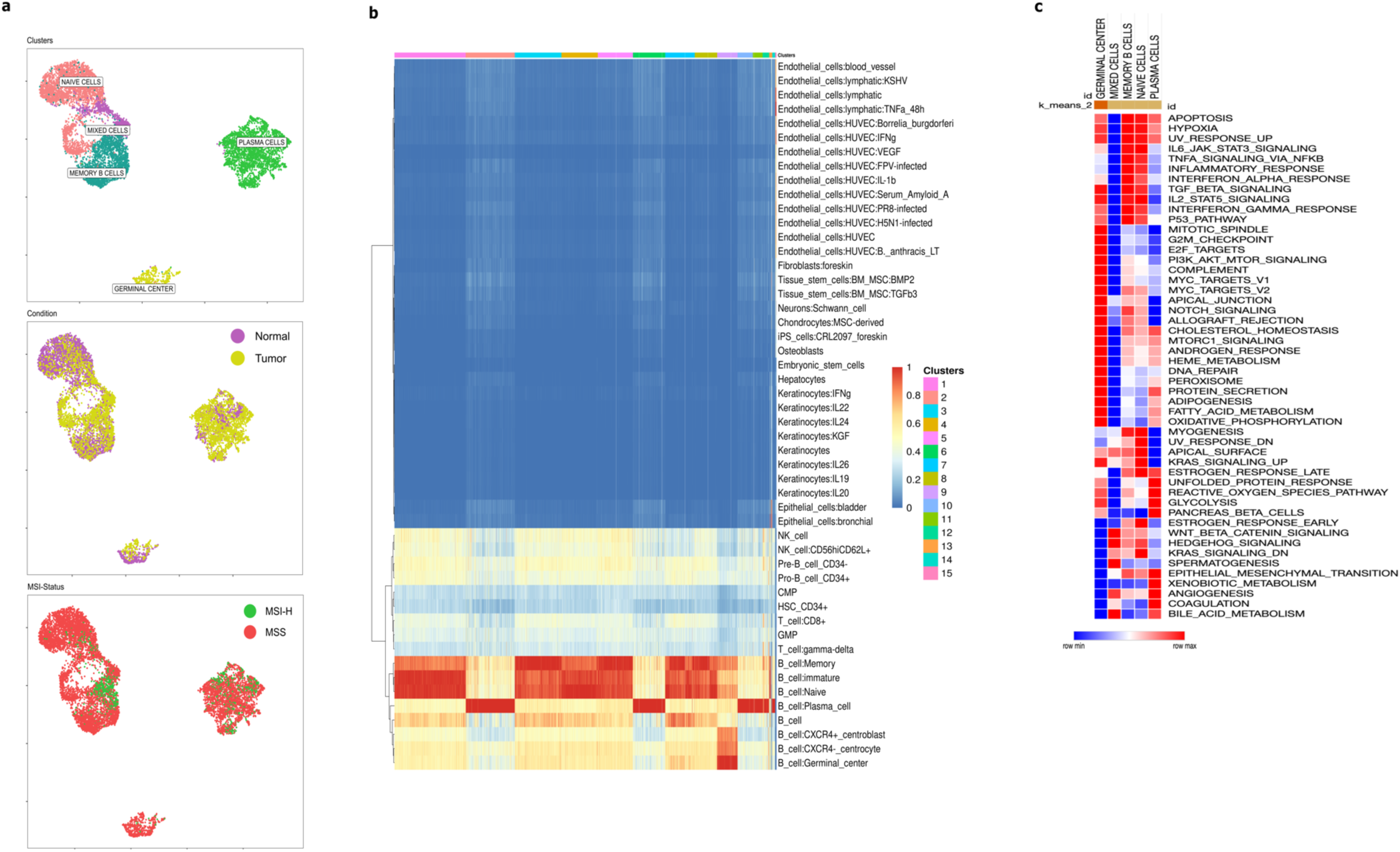
Reclustering of B cells with characterization of clusters and phenotypes. **a**, UMAP depiction of B cell reclustering colored by cluster, malignancy status, and sample of origin. **b**, SingleR heatmap demonstration of B cell distribution within each cluster. **c**, B cell Hallmark pathway analysis by phenotype. scCMS, single-cell consensus molecular subtyping; bulk CMS, consensus molecular subtyping on Bulk RNA-seq data. **Description:** To illustrate characteristics of B cells in CRC we reclustered 9,289 B cells that clearly identified naive cells, memory cells, plasma cells, and germinal center (GC) B cells. All B cell subtypes from the CRC TME and the non-malignant colonic tissue clustered together exhibiting transcriptional similarity among non-tumor and tumor-derived cells. Memory B cells and plasma cells were enriched in tumors, while naive and GC B cells were enriched in nonmalignant tissue.

**Extended Data Figure 3.**
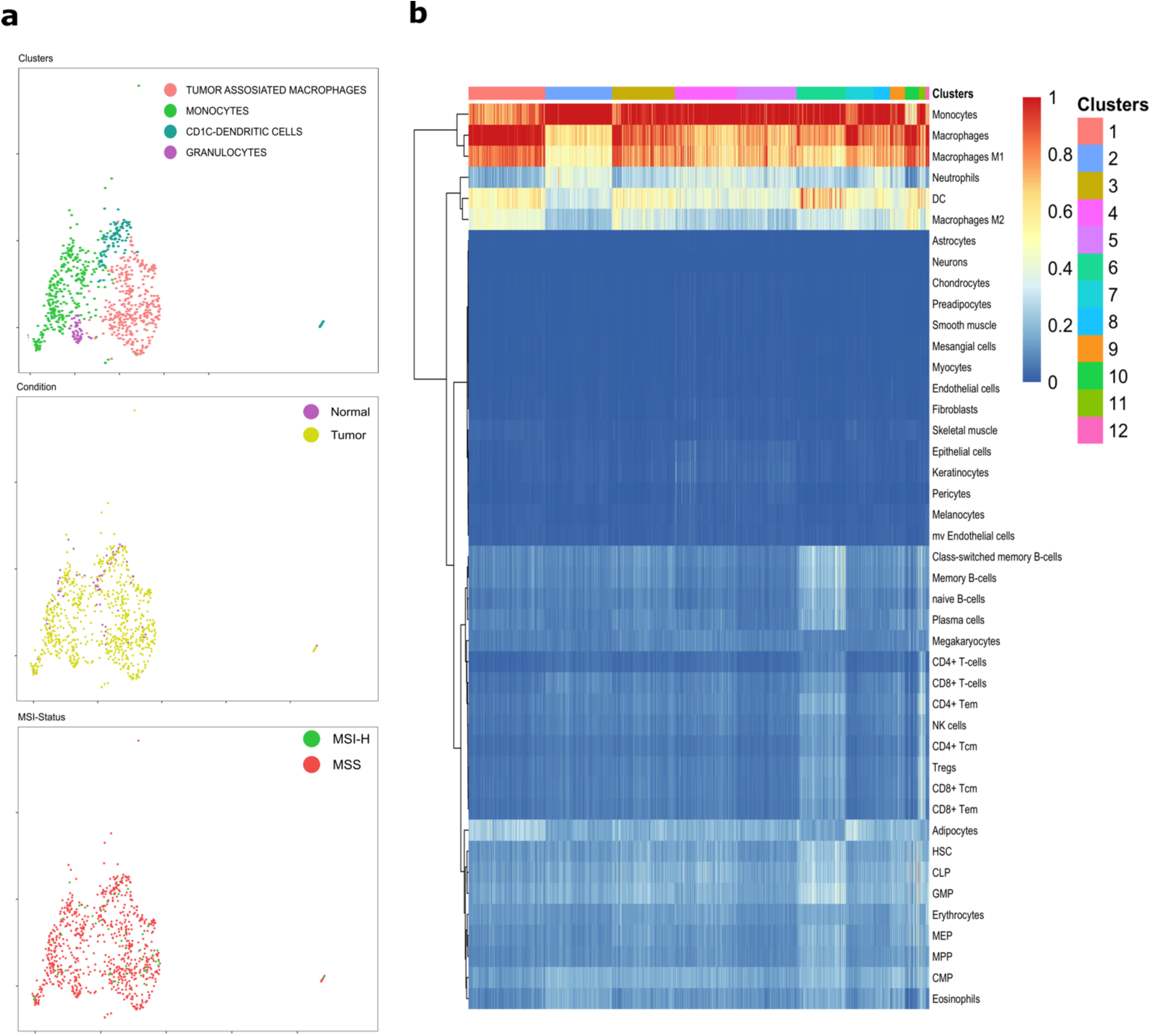
Reclustering of the myeloid cell compartment. **a**, UMAP depiction of myeloid cell reclustering colored by subtype, malignancy status, and sample of origin. **b**, SingleR heatmap demonstration of myeloid cell distribution within each cluster. scCMS, single-cell consensus molecular subtyping; bulk CMS, consensus molecular subtyping on Bulk RNA-seq data. **Description**: We reclustered 819 myeloid cells and identified CD1C+ dendritic cells, tumor associated macrophages (TAM and MRC1+), monocytes (S100A8+), and granulocyte clusters. We recovered key cell types including M2 polarized macrophages, as seen in other tumor types. Monocytes revealed proinflammatory phenotypes (1L1B, S100A8, and S100A9), while TAM showed anti-inflammatory signatures (APOE, SEPP1, and CD163) consistent with the role of TAM in immune suppression and cancer progression (see methods).

